# Pi-Ensemble: Sequence-guided generation of interpolated protein conformational ensembles

**DOI:** 10.64898/2026.08.12.744498

**Authors:** Hassan Nadeem, Diego E. Kleiman, Yuming Zhou, Andrew D. B. Leakey, Diwakar Shukla

## Abstract

Proteins are critical biomolecular machines that populate ensembles of interconverting conformations. Many biological processes depend on transitions between metastable states. Although molecular dynamics (MD) simulations provide a physically grounded route to characterize these motions, routine sampling of large-scale conformational transitions remains computationally demanding. Recent advances in protein structure prediction have created new opportunities for ensemble generation, but many existing approaches require noising inputs, task-specific training, supervised fitting on extensive MD data, or experimentally-informed restraints. Here, we introduce Pi-Ensemble (Predicting Interpolated Ensemble), a sequence-guided framework for generating protein conformational ensembles interpolating between two structural anchor states. Unlike previous methods, Pi-Ensemble alternately leverages inverse-folding and structure-prediction models to propose intermediate conformations between known protein states, generating diverse ensembles without additional training. We evaluate Pi-Ensemble across diverse protein systems, including enzymes, transporters, receptors, and benchmark cases with reference MD simulations or experimental Double Electron-Electron Resonance (DEER) data. Pi-Ensemble recovers physically plausible intermediate conformations, captures transition pathways observed in large-scale MD simulations, and generates structures consistent with experimental distance distributions. Furthermore, Pi-Ensemble-generated conformations provide effective starting seeds for parallel MD simulations, improving conformational exploration and accelerating convergence relative to simulations initiated only from endpoint structures. These results establish sequence-guided structural interpolation as a practical strategy for probing protein conformational landscapes. By generating diverse and physically reasonable conformational proposals without long-timescale MD or model retraining, Pi-Ensemble provides an extensible framework for studying protein flexibility, guiding adaptive sampling, and accelerating mechanistic investigations of protein function.

## 1 Introduction

Protein structure prediction and generation have enabled a wide range of applications in biomolecular modeling, substantially improving our understanding of the structural basis of protein function and design. Recent advances in deep learning-based structure prediction have transformed structural biology by enabling highly accurate prediction of protein structures directly from amino acid sequences.^1–8^ However, proteins are intrinsically dynamic molecules that populate ensembles of conformations rather than single static structures. ^9,10^ Many biological processes, including ligand binding,^11,12^ allostery,^13^ transport,^14,15^ and catalysis,^16,17^ depend on transitions between metastable states and on the thermodynamic and kinetic properties of conformational ensembles. Consequently, there is growing interest in computational methods capable of moving beyond static structure prediction toward the generation and characterization of protein conformational landscapes.^18^

Although early protein structure prediction approaches primarily focused on predicting single stable conformations, numerous recent works have expanded the field to enable prediction of dynamic trajectories, structural ensembles, and residue-level dynamical properties. Existing approaches^19^ for conformational generation and ensemble characterization can be broadly organized into several categories, including physics-inspired coarse-grained machine learning potentials,^20,21^ generative models,^22–32^ input perturbation methods,^33–35^ diffusion-based or score-based guidance methods,^36^ energy-guided sampling,^37^ entropy-guided optimization,^38^ experimentally guided or experimentally constrained prediction,^39^ local transition models,^40–43^ machine learning-based collective variable methods,^44^ distance-restrained prediction,^45^ and predictors of reduced ensemble descriptors.^46–51^ Generative approaches themselves span conceptually distinct paradigms, including AlphaFold2-based generative models,^23,25^ exact likelihood-based models,^27–29^ and methods derived from energy functions.^30–32^ Collectively, these methods provide complementary strategies for exploring protein conformational space. Naturally, each approach has its own advantages and limitations.

In general, the primary advantage claimed by these methods is their ability to generate physically plausible conformational ensembles (or predict ensemble information) without resorting to expensive molecular dynamics (MD) simulations. However, these approaches are often accompanied by significant limitations, including the need for extensive retraining on molecular dynamics trajectories,^25^ reliance on specialized datasets,^48^ or dependence on architecture-specific features.^36^ Other methods require prior knowledge of the system, such as experimental measurements, to correct or bias predicted structures.^39^ Moreover, it should be noted that several of these methods perform best in the regime of structured proteins represented in structural databases, whereas other families of approaches are better suited for intrinsically disordered systems.^52,53^

An interesting concept introduced in the domain of Large Language Models (LLMs) is termed prompt engineering. In natural language settings, researchers have observed that the particular formulation or structure of a query, although semantically identical from a human perspective, may improve or degrade the response of the LLM and even bypass security controls.^54^ In the context of protein structure prediction, one may imagine constructing protein sequences to query desired backbone conformations. This is, in essence, the central goal of inverse folding, a protein design approach that (in most modern iterations) utilizes deep learning models to propose amino acid sequences that satisfy the constraints imposed by an input backbone structure.^55,56^ In this sense, one may envision using inverse folding methods to engineer sequences (the fundamental objects upon which structure prediction models operate) as a means to improve exploration of conformational space. Based on this notion, we introduce Pi-Ensemble (Predicted Interpolated Ensemble), a framework for generating protein conformational ensembles through iterative coupling of inverse folding and structure prediction models. Our approach is distinct from existing input perturbation or guidance methods because it leverages inverse folding models trained on an orthogonal objective (sequence identity recovery) to construct bespoke sequences that prompt structure prediction models to generate conformations that may otherwise remain inaccessible. Importantly, this strategy does not require retraining structure prediction models on ensemble datasets or large-scale MD trajectories.

The conceptual basis of Pi-Ensemble is inspired by iterative peptide binder design pipelines such as BindCraft^57^ or EvoBind,^58^ where peptide sequences are refined such that a structure prediction model infers the formation of a stable complex. In contrast, Pi-Ensemble uses two structural templates (corresponding either to different states of the same protein or to homologous proteins in different poses) as interpolation anchors. Inverse folding models are used to obtain residue identity probability distributions conditioned on each anchor structure, and interpolation is then performed within this probabilistic sequence space. The resulting sequences can differ substantially from the original protein sequence across many positions while still encoding structural compatibility with intermediate conformations.

These generated sequences are subsequently passed through protein structure prediction models to obtain corresponding structures. The predicted conformations are then incorporated into iterative rounds of sequence design and structure prediction, progressively expanding coverage of the conformational landscape between the input anchors. To recover physically realistic representations of the target protein, the original reference sequence is reconstructed onto the generated backbones through side-chain repacking,^59^ followed by energy minimization using classical MD force fields.^60^ A major advantage of this framework is that both the structure prediction and inverse folding components can be used directly in their pretrained forms, enabling straightforward deployment and compatibility with future advances in either model class.

We demonstrate that Pi-Ensemble successfully generates physically plausible conformations that reproduce transition pathways observed in large-scale MD simulations across diverse protein systems. Furthermore, when the generated ensembles are used as seeds for parallel MD simulations, conformational landscapes can be recovered using approximately an order of magnitude less simulation data compared to conventional approaches. Because sampling can be distributed across independently generated structures, the resulting wall-clock acceleration can span several orders of magnitude when sufficient computational resources are available. In this sense, our method can be combined with powerful distributed computing platforms such as Folding@home (F@H)^61^ to substantially accelerate sampling of conformational distributions.

Importantly, Pi-Ensemble produces structurally diverse ensembles that, in some systems, exceed the diversity recovered by existing methods. These results suggest that sequence prompting through inverse folding models enables recovery of previously unseen conformations and that the interaction between inverse folding and structure prediction models produces emergent behavior that cannot be trivially predicted from either component independently. To facilitate adoption and further development, we provide a user-friendly and extensible Python package implementing the Pi-Ensemble framework for efficient generation of protein conformational ensembles.

## 2 Results

In this section, we demonstrate the utility of Pi-Ensemble across a range of applications, from identifying transition states to improving the efficiency of parallelized molecular dynamics simulations. We begin by introducing the Pi-Ensemble framework and summarizing key results. After establishing that Pi-Ensemble generates structurally consistent transition states, we validate its performance against large-scale physics-based simulations.^15,62^ We then show that MD simulations seeded with structures generated by Pi-Ensemble achieve substantially improved efficiency while maintaining or enhancing conformational exploration and convergence metrics. To show the broad scope of the method, we apply Pi-Ensemble to a broader set of 18 protein systems.^63^ Finally we compare the Pi-Ensemble structures with experimental data from Double Electron-Electron Resonance (DEER) spectroscopy.^64^

### 2.1 Pi-Ensemble is a framework for sequence-guided structural interpolation

The primary objective of Pi-Ensemble is to generate plausible protein structures that span the conformational space between two structural anchors. These anchors may correspond to, for example, active and inactive states of a G protein-coupled receptor (GPCR),^65,66^ or inward-and outward-facing conformations of a membrane glucose transporter.^15^ To begin, each anchor structure is independently passed through an inverse folding model, such as ProteinMPNN.^55^ This step produces a conditional probability distribution over amino acid identities at every position in the sequence. We note only probabilistic inverse folding models are compatible with our approach. Sampling from this distribution is expected to yield sequences that are compatible with, and thus fold into, the original input structure. Applying this procedure to both anchors results in two distinct conditional probability distributions. We posit that differences between these distributions encode information about the structural variation between the two anchor states. To leverage this, we construct a linear combination of the distributions using a weighting parameter *λ ∈* [0, 1], which controls the relative contribution of each anchor. The resulting *interpolated* conditional distribution is intended to reflect intermediate sequence preferences associated with conformations between the two endpoints. From this interpolated distribution, we extract a maximum-likelihood sequence, which is then passed to a structure prediction model, such as ESM3,^8^ Boltz,^5,6^ or the more recent ESMFold2^67^ to generate a corresponding protein structure. This workflow is illustrated in Figure 1a. At the end of generation, the native sequence can be backmapped to the backbone structures and sidechains can be repacked using available programs like Rosetta^68^ or MODELLER;^69^ we employ cg2all^59^ for this purpose.

**Figure 1:**
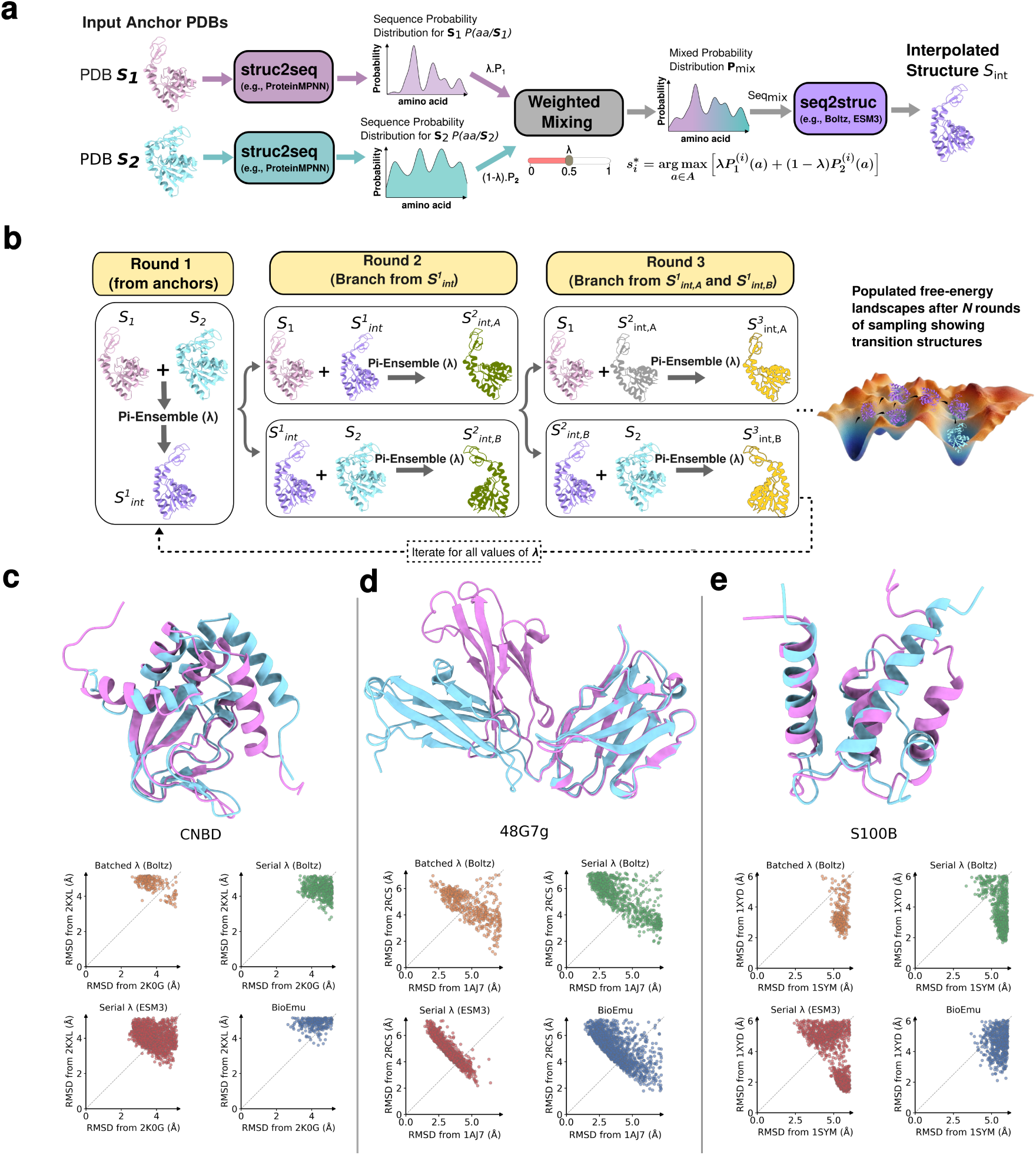
(a) Pi-Ensemble interpolates between anchor structures *S*_1_ and *S*_2_ to span intermediate conformational space. (b) Workflow: each anchor is processed by an inverse folding model; resulting distributions are combined via weighting factor *λ*; the maximum-likelihood sequence is passed to a structure prediction model to generate an intermediate conformation. (c) Iterative application yields an ensemble sampling the underlying free-energy landscape. (d–f) Performance on CNBD, 48G7g, and S100B: top panels show aligned anchors; bottom panels show RMSD projections of generated structures using serial and batch *λ* schemes with ESM3 and Boltz, with BioEmu (blue) for comparison.

After the first round, the anchors are updated and the procedure is applied iteratively to progressively refine the interpolation pathway between the structural templates (Figure 1b). In the first round, the initial two anchors are used to generate the first interpolated structure (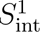). Subsequently in the second round the first anchor (*S*_1_) is held fixed, while the structure predicted in the previous round (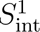) replaces the second anchor (*S*_2_). Alternatively, *S*_2_ is held fixed, and 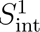 replaces the first anchor. In both cases, the method iteratively updates one endpoint with the most recently predicted intermediate, effectively pushing the structural trajectory toward either of the original anchors. This scheme can be viewed as first generating an initial intermediate structure, and then progressively refining the conformational pathway through successive interpolation steps. Importantly, the choice of *λ* at each step influences the relative proximity of the predicted structure to either anchor, thereby controlling the direction and extent of progression along the transition. Consequently, multiple iterative rounds, combined with varying *λ* values at each step, produce a diverse ensemble of structures that more fully spans the conformational space between the original anchor states. The next section explains how the interpolation is performed in detail.

### 2.2 Proposed interpolation schedules enable exhaustive sequence generation

A central implementation choice in Pi-Ensemble is how to schedule the interpolation parameter *λ*. Initially, two inverse-folding probability distributions, *P*_1_ and *P*_2_, are obtained from the two anchor structures after mapping both templates onto a common residue indexing scheme. This alignment step is important when the two templates contain missing residues or differ slightly in sequence coverage, because interpolation must be performed between probability distributions defined over the same set of positions. This step also allows to input two homologous (or even unrelated) structures as anchors, as long as residue positions are suitably aligned. For each residue position *i* and amino acid identity *a*, Pi-Ensemble defines an interpolated distribution

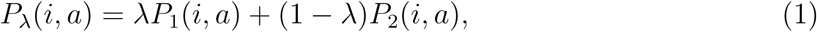

where *λ ∈* [0, 1] controls the contribution of each anchor. The sequence used to query the structure prediction model is obtained by maximum-likelihood decoding,

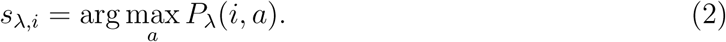

Thus, *λ* does not directly interpolate Cartesian coordinates. Instead, it interpolates structural preferences encoded in the inverse-folding probability distributions, and the structure prediction model maps the resulting sequence to a three-dimensional conformation.

We implemented two complementary strategies for choosing *λ*: a serial *λ* schedule and a batch *λ* schedule. The serial schedule uses a fixed grid of user-defined *λ* values, whereas the batch schedule adaptively identifies the values of *λ* at which the decoded maximum-likelihood sequence changes. These two strategies differ in their computational profile and in the type of conformational diversity they are designed to capture.

#### 2.2.1 Serial interpolation

In the serial *λ* algorithm, the user defines a fixed set of mixing weights using a starting value, ending value, and uniform step size. For each value of *λ*, interpolation is performed in both “directions.” First, *S*_1_ is held fixed as the anchor and the second template is treated as the mobile endpoint. In the other direction, the roles of the two templates are reversed. At each iteration, the anchor and mobile inverse-folding distributions are mixed according to the predefined *λ*, the maximum-likelihood interpolated sequence is decoded, and the resulting sequence is passed to the structure prediction model. The newly predicted structure is then processed again by the inverse-folding model, producing a new probability distribution that replaces the previous mobile endpoint in the next round. In this way, the serial algorithm constructs an iterative trajectory in probability-distribution space: one endpoint remains fixed, while the other endpoint is repeatedly updated using the most recently generated structure.

This recursive update is intended to progressively move through sequence prompts that are compatible with structures between the original anchors. Running the procedure in both directions reduces dependence on the arbitrary choice of which anchor is fixed and allows the algorithm to explore paths biased toward either endpoint. The fixed *λ* grid also provides direct control over the granularity of the interpolation. Small step sizes generate many closely related sequence prompts, while larger step sizes provide a coarser but less expensive exploration.

#### 2.2.2 Batched interpolation

The batch *λ* algorithm uses a different scheduling principle. Rather than evaluating a uniform grid of *λ* values, it exploits the fact that the decoded sequence changes only when the identity with maximum probability changes at one or more residue positions. For a given pair of anchor distributions, the algorithm computes the critical values of *λ* at which two amino acid probabilities become equal at the same residue position:‘

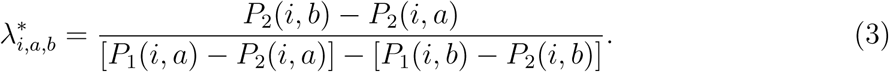

Conceptually, a value is considered critical when the probability of one residue identity (*b*) exceeds the probability of another residue identity (*a*) as *λ* crosses that value. This can be read as: at 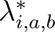, position *i* switches from residue *a* to residue *b*. Only finite critical values satisfying 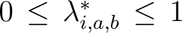 are retained, ensuring that the interpolation remains within the range bounded by the two anchor distributions. These critical values define intervals over which the maximum-likelihood decoded sequence is constant. The algorithm then samples representative values from these intervals, using midpoints between consecutive critical values and including the endpoints when appropriate. This produces a nonuniform, sequence-aware schedule of *λ* values.

For each selected *λ*, the batch algorithm decodes the corresponding maximum-likelihood sequence and removes redundant sequences using a minimum edit-distance criterion set by the user. Under default parameters, all unique sequences are included. This optional filtering step avoids spending structure-prediction evaluations on sequence prompts that differ only slightly from previously selected prompts. The filtered sequences are then submitted to the structure prediction model, using batched prediction when available. Each predicted structure is subsequently passed back through the inverse-folding model to obtain a new probability distribution, which can be used in later rounds.

The iterative anchor-selection scheme in the batch algorithm is designed to expand the interpolation pathway without requiring a fixed *λ* grid. In the first round, interpolation is performed directly between the two original templates. In the second round, the generated sequence with the largest edit distance is selected as a common intermediate anchor and paired separately with each original template. In subsequent rounds, the algorithm separately tracks structures generated in each direction and selects, for each branch, the structure with the largest edit distance from the previous round as the new mobile endpoint. This strategy encourages the algorithm to propagate toward sequence regions that are most distinct from the current anchor, while retaining the original templates as stable reference endpoints.

The serial and batch schedules therefore represent two related but distinct approaches to interpolation. The serial schedule is simple, deterministic, and easy to control through the number of rounds and the spacing of the *λ* grid. It is useful when dense sampling of the interpolation path is desired, but it may repeatedly evaluate *λ* values that decode to identical or very similar sequences. In contrast, the batch schedule is adaptive: it focuses computation on values of *λ* that produce distinct maximum-likelihood sequences and therefore distinct prompts to the structure prediction model. This can substantially reduce redundant evaluations and allows the number of generated structures to depend on the information content of the two inverse-folding distributions (more interpolated sequences are queried for very disparate templates or for larger proteins). In practice, both strategies provide useful conformational ensembles, and their complementary behavior allows Pi-Ensemble to balance exhaustive interpolation with computational efficiency.

### 2.3 Interpolated ensembles span the conformational space between anchors

As an initial test of performance, we apply the Pi-Ensemble framework to three protein systems (Figure 1c–e, top panels): the bacterial cyclic nucleotide-activated K^+^ channel binding domain (CNBD), immunoglobin 48G7 germline (48G7g), and the calcium-binding protein S100B. We evaluate three variants of Pi-Ensemble: batch *λ* using Boltz as the structure generator (orange), serial *λ* with Boltz (green), and serial *λ* with ESM3 (red). For reference, we also include results from BioEmu^25^ (blue). To assess whether Pi-Ensemble generates transition structures between the two anchors, we compute the root mean square deviation (RMSD) of each structure with respect to both anchors and visualize the results (Figure 1c–e, lower panels). In these plots, each axis represents the RMSD to one of the anchors. To enforce a strict criterion of structural soundness, we discard any structure whose RMSD to either anchor exceeds the RMSD between the two anchors themselves. We note that this filtering is conservative, as valid intermediate structures may violate this criterion. Across all three systems, Pi-Ensemble successfully generates intermediate structures. The 48G7g system shows the most complete coverage of the RMSD space, with structures distributed near both anchors as well as a substantial population in between. In contrast, all methods exhibit difficulty for S100B, where structures fail to approach the 1SYM anchor.

An interesting observation is that all three Pi-Ensemble variants exhibit distributional trends that closely resemble those produced by BioEmu. This similarity is notable because BioEmu was trained on hundreds of milliseconds of MD simulations in addition to millions of protein structures, whereas ESM3 was not trained on MD data and Boltz-2 was trained on smaller MD datasets (both models were fitted on comparably large structural datasets). Despite these differences in training data, the resulting conformational distributions are remarkably similar. Based on these results, we speculate that the ability to recover conformational landscapes from structural anchors without model retraining will become an attractive alternative for systems that are not represented in MD datasets. Unfiltered RMSD scatter plots for the three systems are shown later in Figure 5.

### 2.4 Pi-Ensemble captures conformational pathways observed in large-scale molecular dynamics simulations

Physics-based MD simulations have long served as a central approach for probing conformational heterogeneity in protein systems.^70^ By explicitly modeling atomic interactions over time, MD provides a physically grounded description of protein motions and enables detailed characterization of structural ensembles and transition pathways. However, these simulations are computationally demanding, owing to the need for small integration time steps and repeated evaluation of complex force fields. As a result, accessing long timescales or rare conformational transitions remains challenging in many cases. Despite these limitations, MD has provided critical insights into protein dynamics and continues to serve as an important reference for understanding conformational landscapes. Given this role, it is natural to validate Pi-Ensemble generated structures against MD simulations. In particular, we aim to assess two complementary aspects. First, we evaluate how many generated structures are *close* to those sampled in MD simulations, which corresponds to the precision of our framework. Second, we assess whether the generated ensemble adequately covers a substantial portion of the conformational landscape explored by MD, which corresponds to recall. The first criterion measures how physically realistic the generated structures are. Not only in terms of avoiding steric clashes or non-physical geometries, but also in their ability to resemble conformations that arise in physics-based simulations. The second criterion is more stringent, as it tests whether Pi-Ensemble can produce a sufficiently diverse ensemble to span broad regions of the conformational landscape, including metastable states and transition pathways.

To this end, we use an extensive MD simulation dataset from Weigle *et al.* (2024),^15^ who investigated AtSWEET13, an *Arabidopsis thaliana* sugar transporter responsible for mediating sugar transport across cellular membranes.^71^ For our analysis, we focus on simulations of the *apo* (ligand-free) state of AtSWEET13, comprising a cumulative simulation time of 132 *µ*s. Figure 2a shows the inward (pink) and outward (teal) facing conformations identified from these simulations. Gating residues are colored in red. These structures are used as input anchors for Pi-Ensemble to generate interpolated conformations, as shown in Figure 2b. Pi-Ensemble performs well in this setting, generating intermediate conformations that smoothly span the RMSD space between the two anchors. Even under a strict cutoff on inter-anchor RMSD (4.26 ÅA), a substantial number of structures populate the intermediate region, while also clustering within 2 ÅA of each anchor. We further examine the kernel density estimate (KDE) of predicted Local Distance Difference Test (pLDDT) scores in Figure 2c, which shows that the ensemble is dominated by high-confidence structures. To quantify agreement with reference MD simulations, we compute the RMSD of each Pi-Ensemble-generated structure to its closest frame in the MD trajectories. The resulting distribution, shown in Figure 2e, peaks at 1.5 ÅA, indicating strong structural consistency with MD-sampled conformations. While RMSD-space coverage is encouraging (Figure 2b), a more informative assessment considers sampling along reaction coordinates that drive the conformational transition. In the original AtSWEET13 study, gating residues were identified and used to define intracellular (IC) and extracellular (EC) gating distances. We therefore project Pi-Ensemble-generated structures onto this two-dimensional coordinate space in Figure 2d. For comparison, we also overlay the free-energy landscape obtained from MD simulations. We observe that most generated structures fall within the regions of the free-energy landscape and span a wide range of IC and EC gating distances. Beyond the inward-and outward-facing states, Pi-Ensemble also captures occluded and intermediate transition conformations. This indicates that Pi-Ensemble not only generates physically plausible structures, but also explores the underlying free-energy landscape without any *a priori* knowledge of the reaction coordinates, here defined by the gating distances. These results were obtained using batch *λ* with Boltz as the structure generator. Serial *λ* with Boltz yields comparable results (Figure S1). In contrast, BioEmu and other variants of Pi-Ensemble show reduced coverage of the free-energy landscape, with a pronounced bias toward inward-facing conformations near the crystallographic state (PDB ID: 5XPD), see Figure S1 and Figure S2.

**Figure 2:**
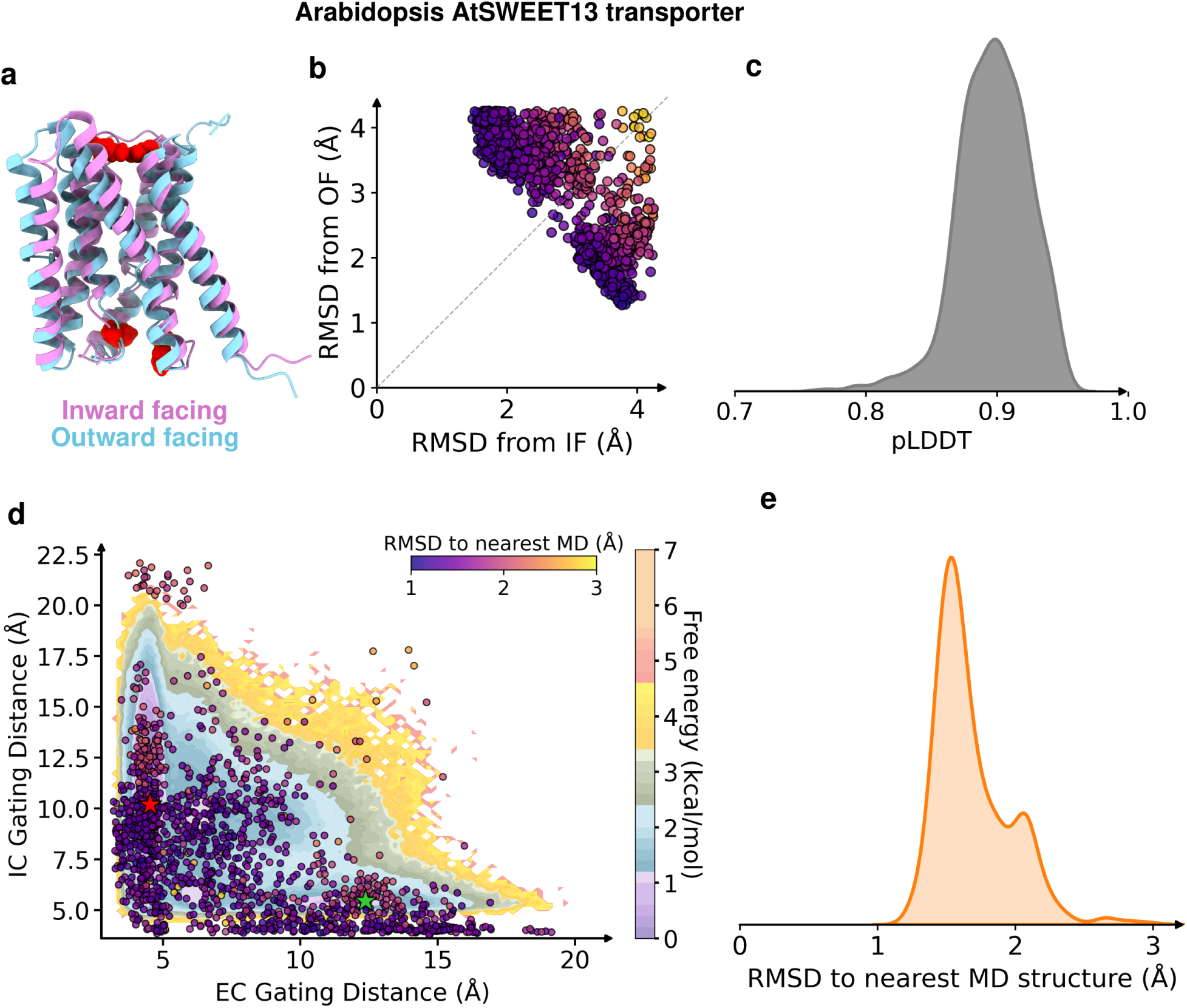
(a) Inward-and outward-facing structures of the *A. thaliana* SWEET13 sugar transporter. (b) Pi-Ensemble generated structures projected onto RMSD space relative to the two anchor structures. (c) Kernel density estimate (KDE) of the predicted Local Distance Difference Test (pLDDT) scores for structures generated from interpolated sequences. (d) Projection of Pi-Ensemble generated structures onto the two-dimensional space defined by intra-and extracellular gating distances, overlaid on the *ground-truth* free energy landscape obtained from 132 *µ*s MD simulations reported by Weigle *et al.* (2024).^15^ (e) KDE of the RMSD between each Pi-Ensemble generated structure and its nearest neighbor in the MD simulation ensemble.

### 2.5 Pi-Ensemble seeds accelerate conformational sampling

In the previous section, we showed that Pi-Ensemble generated structures are structurally valid and collectively span the free-energy landscape of a complex biomolecular system sampled over hundreds of microseconds of physics-based MD simulations. In general, performing simulations on timescales ranging from microseconds to milliseconds remains computationally intractable without access to specialized computational infrastructure such as distributed computing platforms.^61^ This presents a significant challenge, as many biologically important processes, including domain rearrangements, allosteric transitions, and other large-scale conformational changes, occur on timescales of microseconds and beyond. Consequently, directly observing such events through conventional MD simulations is often prohibitively expensive. Traditionally, MD simulations are initialized from structures determined using experimental techniques such as X-ray crystallography,^72^ Nuclear Magnetic Resonance (NMR) spectroscopy,^73^ and cryo-electron microscopy (cryo-EM).^74^ More recently, advances in modern machine learning (ML) architectures have substantially improved the fidelity and reliability of structure prediction models. As a result, a natural extension is to initialize simulations from ML-generated conformations^75–77^ rather than relying exclusively on experimentally resolved structures. This is particularly attractive in the context of conformationally heterogeneous systems, where ML-generated ensembles may provide access to structurally diverse regions of the underlying energy landscape that would otherwise require extensive sampling to discover. Given *n* valid starting conformations, two distinct forms of acceleration can be expected. First, compared to initiating simulations from a single structure, distributing simulations across *n* independent seeds can reduce the effective wall-clock time by up to a factor of *n*, assuming sufficient parallel computational resources are available. This parallelization strategy enables simultaneous exploration of multiple regions of conformational space. The second form of acceleration arises in the context of sampling rare events. Observing a rare event that requires crossing a free-energy barrier depends primarily on the total accumulated simulation time rather than the duration of any individual simulation. This is because the probability of overcoming a free-energy barrier is governed by the total number of crossing attempts made across all trajectories.^78^ Importantly, these *n* simulations evolve independently and therefore will not explore phase space identically to a single continuous trajectory that is *n* times longer. However, adaptive sampling^79–82^ provides a practical framework for generating such independent parallel trajectories by iteratively launching new simulations from underexplored or kinetically relevant conformations. One can then use coupling schemes and statistical frameworks such as Markov state models (MSMs),^83^ which integrate information across independent trajectories to reconstruct long-timescale kinetics and thermodynamic properties.

To investigate whether Pi-Ensemble generated structures can serve as viable starting seeds, we present a Pi-Ensemble augmented framework for MD simulations, depicted in Figure 3a. Using two available anchor structures, we employ Pi-Ensemble to generate an ensemble of diverse protein conformations. This ensemble can then be clustered, with cluster representatives selected for seeding short parallel simulations. For systems with known important features or well-characterized collective variables, clustering can be performed in the corresponding feature space. However, for our two test cases, we perform clustering in RMSD space to represent an agnostic scenario in which no prior physical insight into the system is assumed.

**Figure 3:**
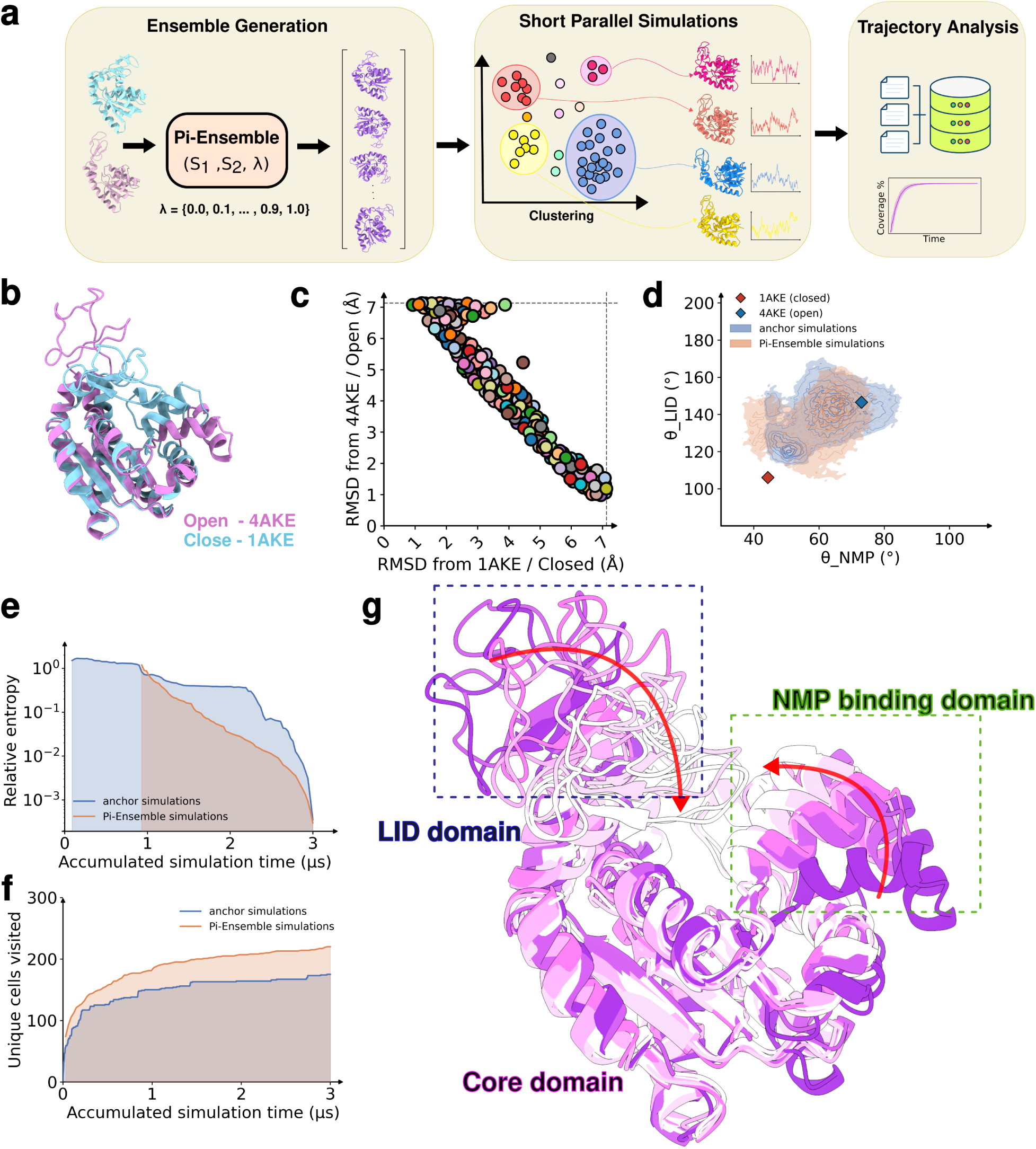
(a) Workflow using Pi-Ensemble to generate seed structures for parallel simulations. (b) Open (PDB ID: 4AKE) and closed (PDB ID: 1AKE) AdK states. (c) RMSD-to-anchor projection of 298 clustered Pi-Ensemble structures used as simulation seeds. (d) MD results from anchor-only (blue) versus anchor plus Pi-Ensemble seeds (orange); anchors shown as diamonds and simulations projected onto lid and NMP domain angles. (e) Relative entropy and (f) conformational exploration comparing anchor-only (blue) and Pi-Ensemble-initiated simulations. (g) Representative Pi-Ensemble structures showing transition from open (purple) to closed (white), with colors shifting from dark (open) to light (closed).

We consider adenosine kinase (AdK) as our first test case. AdK is an enzyme that phosphorylates adenosine to AMP, thereby regulating extracellular adenosine levels.^84^ It plays a key role in purine metabolism and has been implicated in epilepsy, pain, and neurological disorders.^85^ We use two publicly available open (PDB ID: 4AKE)^86^ and closed (PDB ID: 1AKE)^87^ structures as input anchors, Figure 3b. Any ligands and non-protein atoms were removed. Figure 3c shows the RMSD space projection of the 298 cluster representatives used for simulations. These clusters were derived by combining structures generated from Pi-Ensemble with Boltz and ESM3 (Figure S3). We ran two sets of MD simulations. The first consisted of two independent long simulations initiated from the two anchor structures, for a total of 3 *µ*s (2*×*1.5 *µ*s). For the Pi-Ensemble seeds, we ran a comparable total of 3 *µ*s using 298 seeds plus the 2 anchor structures (300*×*10 ns). For analyzing simulation performance, we use the well-characterized angles between the core domain and both the lid and NMP domains, referred to as the lid and NMP angles, respectively (Figure 3d). We observe that both sets of simulations eventually sample a very similar conformational space projected onto the lid and NMP angles, owing to the relatively low barrier between the open and closed conformations in this system. To quantify which set of simulations explored the space more rapidly, we divided the NMP and lid angle space into a (20, 30) grid and counted the number of unique cells visited by each method (Figure 3f). We observe that the Pi-Ensemble simulations consistently visit a greater number of cells at increasing simulation times, demonstrating that a multi-seed approach outperforms traditional sampling even in a system with relatively small barriers. We also investigated the thermodynamic convergence of the two simulation sets. We constructed MSMs independently for each set and determined their converged transition probability matrices. At increasing intervals of accumulated simulation time, we generated MSMs for both systems and computed the relative entropy (see Methods) between the sampled transition probability matrix and the converged transition probability matrix for that system, Figure 3e. We observe that the Pi-Ensemble simulations approach the converged transition probability distribution significantly faster than simulations initiated solely from the anchor structures. Using representative frames, Figure 3g, we show that Pi-Ensemble captures a gradual and systematic transition between the closed and open conformations with respect to both the LID and NMP domain movements. The details of simulation parameters and MSM construction are described in the Methods section.

Next, we apply the Pi-Ensemble augmented MD simulation pipeline to glutamine-binding protein (GlnBP). GlnBP is a periplasmic protein in *Escherichia coli* that captures L-glutamine and delivers it to a membrane transporter, serving as the entry point for active amino acid uptake.^88^ It adopts a bilobal architecture that transitions between open and closed conformations through inter-domain hinge movements and a network of stabilizing interactions around the bound ligand. To run Pi-Ensemble, we use the two publicly available open (PDB ID: 1GGG) and closed (PDB ID: 1WDN) structures as input anchors, Figure 4a. We then perform clustering and select 186 representative structures as seeds for Pi-Ensemble simulations, Figure 4b. As before, we simulate the anchor and Pi-Ensemble systems for an equal total simulation time of 3.76 *µ*s: 2*×*1.88 *µ*s for the anchor simulations and 188*×*20 ns for the Pi-Ensemble simulations. We featurized the simulation trajectories using two conformational order parameters that distinguish the open and closed states of GlnBP: (i) a hinge-bending angle defined by the centers of mass of the large domain, linker/hinge domain, and small domain, *θ*, and (ii) the center-of-mass distance between the large and small domains, *d*. Figure 4c shows the density map of both simulation sets projected onto these features. We observe that Pi-Ensemble simulations successfully sample a broad conformational space spanned by these features, including regions around the anchors as well as the transition region between them. In contrast, the anchor simulations appear to remain concentrated around the 1GGG (open) conformation. To investigate this further, we plot the anchor simulations as a scatter plot in Figure 4f. We observe that the trajectory initiated from the open conformation (dark blue) samples around the same state throughout the simulation, whereas the trajectory initiated from the closed conformation transitions toward the open state and remains there. This demonstrates that the anchor simulations not only fail to sufficiently sample both states, but also provide too few transitions between them to adequately span the transition region. Examining the marginal distributions for the inter-domain distance and hinge-bending angle in Figure 4d, we observe that Pi-Ensemble simulations sample a broader range of both features, while the anchor simulations sample a much narrower region centered around the open conformation. As before, we quantify conformational exploration in Figure 4g, where Pi-Ensemble simulations clearly outperform the anchor simulations in terms of the diversity of sampled conformations. Figure 4h shows representative structures from the Pi-Ensemble simulations, exhibiting a systematic conformational shift from open (dark) to closed (light) states. We also compared the Pi-Ensemble simulations with a recently deposited simulation dataset,^62^ which accumulated a total simulation time of 53.6 *µ*s. Figure 4g shows this comparison. It can be clearly observed that Pi-Ensemble simulations closely match the conformational sampling achieved by the larger dataset, while providing significant improvement in the transition region. Importantly, the total amount of simulation data generated by Pi-Ensemble was approximately fourteen-fold smaller. Together, these observations demonstrate that Pi-Ensemble structures serve as suitable seeds for simulations and, due to their structural validity and conformational diversity, provide orders-of-magnitude improvements in both wall-time efficiency and conformational sampling.

**Figure 4:**
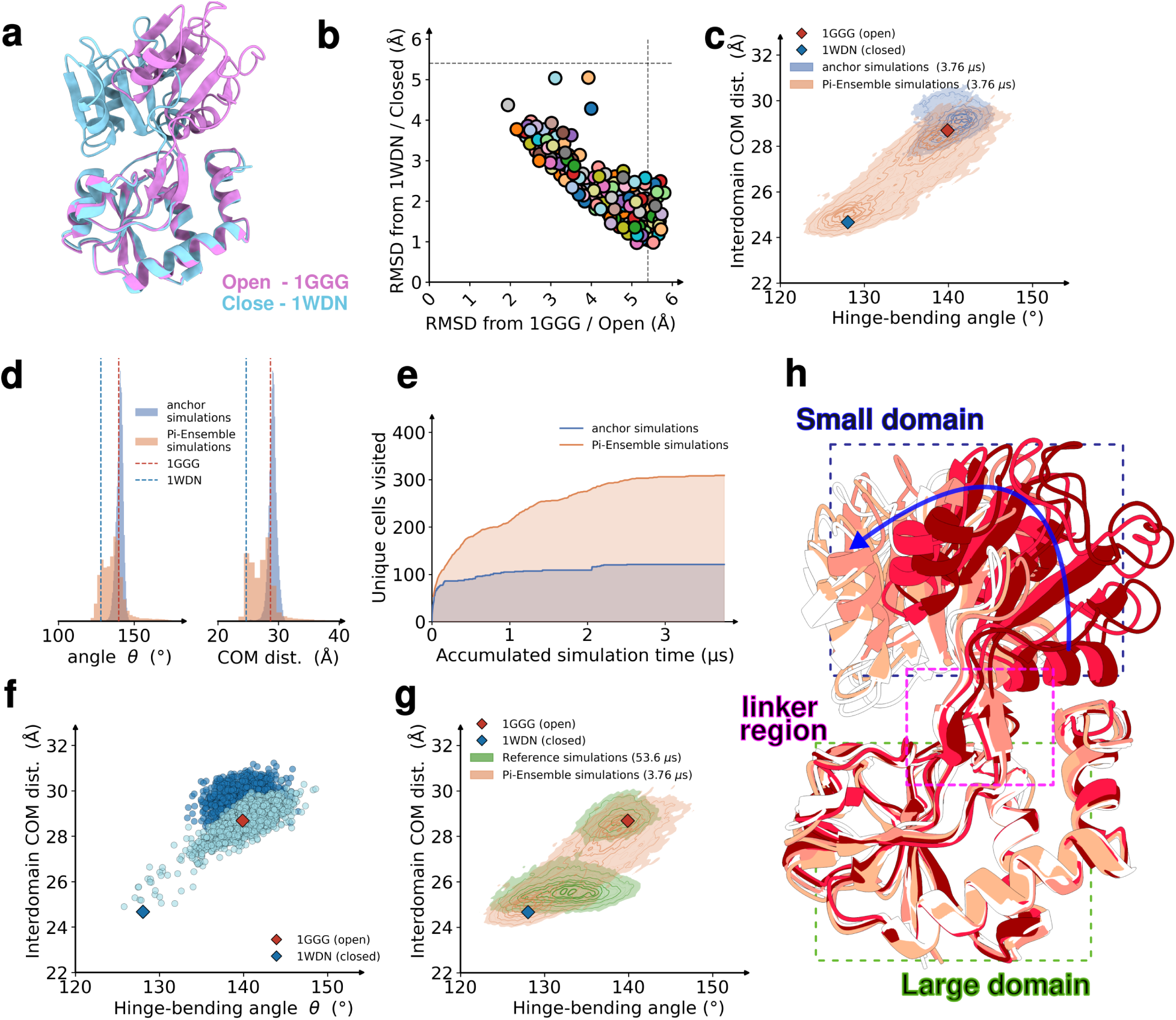
(a) Open (PDB ID: 1GGG) and closed (PDB ID: 1WDN) states of the glutamine-binding protein (GlnBP). (b) RMSD-to-anchor projection of 186 clustered structures generated from Pi-Ensemble outputs using regular clustering; these structures were used as initial seeds for subsequent simulations. (c) Comparison of MD simulations initialized from anchor structures only (blue) versus those initialized from both anchors and Pi-Ensemble structures (orange). Anchor structures are shown as diamonds. Simulations are projected onto the hinge-bending angle and the interdomain center-of-mass distance. (d) Marginal distributions of the hinge-bending angle and interdomain center-of-mass distance for both anchor-only and Pi-Ensemble simulations. (e) Comparison of conformational exploration between anchor-only (blue) and Pi-Ensemble-initiated simulations. (f) Scatter plot of anchor-only simulations. (g) Overlay of Pi-Ensemble simulations with reference simulations^62^ totaling 53.6 *µ*s. (h) Representative Pi-Ensemble structures illustrating a transition from open (red) to closed (white) states, with colors gradually shifting from dark (open) to light (closed). experimental structures also contained unresolved residues, which were reconstructed using MODELLER.^69^

### 2.6 Pi-Ensemble generalizes across diverse protein systems

To demonstrate the utility of Pi-Ensemble across a broad range of protein systems, we utilize a benchmark set^63^ consisting of 18 proteins spanning multiple functional and structural classes. Supporting Table 1 summarizes the details of these systems. The dataset includes kinases, ion transporters, ion-binding proteins, chaperones, signaling proteins, and enzymes, among others, and was selected to encompass a wide range of conformational behaviors. Protein sizes vary substantially, ranging from S100B (83 residues) to HSPD1 (527 residues). The associated conformational transitions are similarly diverse, including domain rotations, hinge-bending motions, helix rearrangements, and combinations of multiple collective motions. In addition, the RMSD between anchor structures spans a broad range, from 3.32 ÅA to 13.97 ÅA, reflecting varying magnitudes of structural change across the benchmark. Several

Pi-Ensemble successfully generated interpolated conformations for the majority of benchmark systems, as shown in Figure 5. For most proteins, the generated structures densely populated the conformational region between the two anchor states when projected onto RMSD coordinates relative to each anchor. Rather than clustering tightly around one endpoint, the ensembles generally formed continuous distributions spanning the transition landscape, indicating that Pi-Ensemble is capable of sampling structurally distinct intermediate states between experimentally resolved conformations. In many systems, these intermediates appeared smoothly distributed along the interpolation pathway, suggesting that the framework captures gradual conformational progression rather than producing isolated or discontinuous structural snapshots. Importantly, Pi-Ensemble was able to generate meaningful intermediates even for systems undergoing relatively large conformational changes. Proteins with substantial domain rearrangements or collective motions still exhibited broad coverage of the interpolated conformational space, demonstrating that the framework is not restricted to small near-native perturbations. A small subset of systems, such as S100A1, produced structures with relatively high RMSD relative to both anchor conformations. These outlier structures may correspond to partially unfolded or otherwise nonphysical conformations, likely arising from limitations in the underlying structure prediction models or from the sequence evolution diverging too far to fold near the native fold.

**Figure 5:**
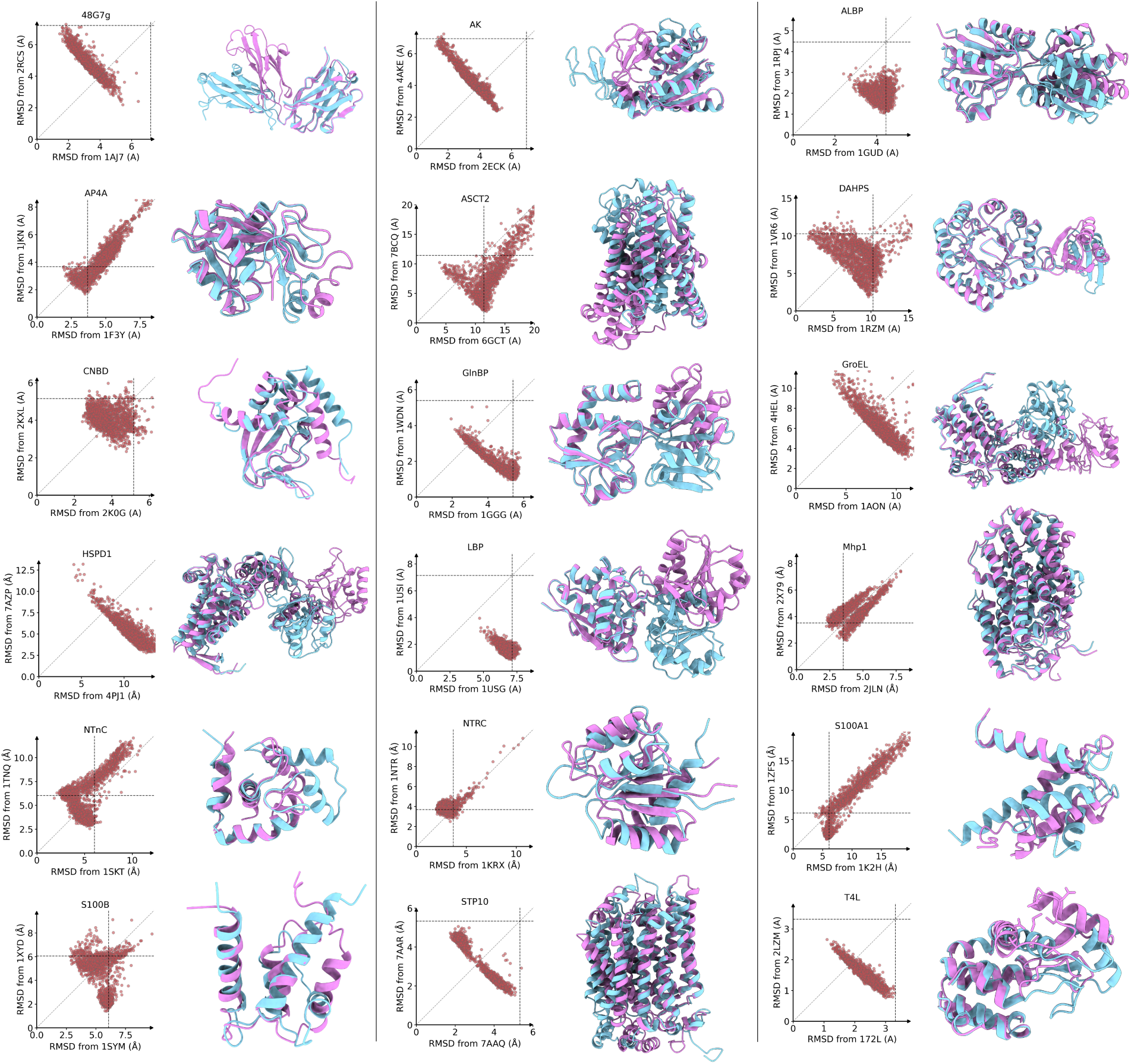
Set of 18 proteins illustrating the performance of Pi-Ensemble with serial *λ* and ESM3 as the structure generator. For each protein, the right panel shows the alternate conformations used as anchor structures (teal and pink), while the left panel presents the RMSD-to-anchor projection of the Pi-Ensemble generated structures.

For a more quantitative assessment, we evaluated the generated structures using two complementary metrics. First, we quantified the proportion of structures that satisfied the RMSD filtering criterion, where the filtering threshold was defined by the RMSD between the two anchor conformations (Table 1). This metric provides a measure of the structural plausibility and validity of the generated ensembles. Second, we calculated the mean RMSD-axis coverage to assess how effectively the generated structures populated the conformational space between the anchor states. To do so, the RMSD-to-anchor coordinate was partitioned into 20 equally spaced bins, and the fraction of populated bins was computed (Table 2). Higher coverage indicates a smoother and more continuous transition between the two end-point conformations. Overall, the Pi-Ensemble variants exhibited strong performance across the benchmark systems. Among them, the Pi-Ensemble implementation using ESM3 as the structure generator produced the highest proportion of structures passing the RMSD filter. RMSD-axis coverage varied across protein systems and model variants, suggesting that the choice of structure generator subtly influences the conformational diversity and distribution of the generated ensembles. Results obtained with BioEmu are included as a reference for comparison. Overall, these results demonstrate that Pi-Ensemble can robustly generate structurally diverse interpolated ensembles across proteins spanning a wide range of sizes, folds, and conformational transition types. The benchmark highlights the ability of the framework to recover continuous intermediate conformational landscapes without requiring explicit MD simulations, transition path sampling, or additional system-specific training. We compare results obtained using different structure prediction models, ESM3 and ESM-Fold2, as well as different interpolation schedules, serial and batch, in Figures S8–S12. For reference, BioEmu results for all 18 proteins are included in the same comparison.

**Table 1:** Structures passing the RMSD filter (%).

| Protein | Pi-Ensemble |  |  | BioEmu |
| --- | --- | --- | --- | --- |
|  | Pi-ESM3 (Serial) | Pi-ESMFold2 (Serial) | Pi-ESMFold2 (Batch) |  |
| 48G7g | **100.0** | 76.5 | **100.0** | 83.4 |
| AK | **99.8** | 99.7 | 98.5 | 85.8 |
| ALBP | **80.2** | 27.9 | 25.8 | 61.9 |
| AP4A | 17.6 | **60.3** | 51.0 | 14.6 |
| ASCT2 | 33.2 | 23.5 | 70.8 | **97.1** |
| CNBD | **94.6** | 45.2 | 72.9 | 15.2 |
| DAHPS | 78.4 | 37.1 | 33.6 | **88.0** |
| GlnBP | **90.0** | 53.8 | 69.7 | 83.2 |
| GroEL | **98.7** | 88.9 | 87.5 | 94.2 |
| HSPD1 | **99.7** | 69.0 | 96.9 | 87.6 |
| LBP | 87.1 | 92.0 | **94.5** | 78.9 |
| Mhp1 | 2.8 | 1.5 | 9.1 | **60.2** |
| NTRC | 27.2 | 28.4 | **28.9** | 1.7 |
| NTnC | **51.6** | 46.5 | 44.5 | 16.9 |
| S100A1 | **33.9** | 12.2 | 9.2 | 25.5 |
| S100B | **67.5** | 48.9 | 50.5 | 32.0 |
| STP10 | 98.1 | **100.0** | **100.0** | 94.0 |
| T4L | 98.1 | 96.2 | **100.0** | 81.9 |

**Table 2:** Mean RMSD-axis coverage after filtering (20 bins; %).

| Protein | Pi-Ensemble |  |  | BioEmu |
| --- | --- | --- | --- | --- |
|  | Pi-ESM3 (Serial) | Pi-ESMFold2 (Serial) | Pi-ESMFold2 (Batch) |  |
| 48G7g | 67.5 | 70.0 | **85.0** | 82.5 |
| AK | 67.5 | **80.0** | 75.0 | **80.0** |
| ALBP | 52.5 | **77.5** | **77.5** | **77.5** |
| AP4A | **52.5** | **52.5** | 50.0 | 45.0 |
| ASCT2 | **77.5** | 57.5 | 57.5 | 75.0 |
| CNBD | 55.0 | **60.0** | 57.5 | 42.5 |
| DAHPS | 85.0 | 80.0 | 70.0 | **87.5** |
| GlnBP | 70.0 | **82.5** | **82.5** | **82.5** |
| GroEL | 77.5 | 77.5 | **87.5** | 85.0 |
| HSPD1 | 77.5 | 80.0 | 82.5 | **87.5** |
| LBP | 40.0 | **90.0** | 77.5 | 80.0 |
| Mhp1 | 37.5 | 30.0 | 40.0 | **47.5** |
| NTRC | **37.5** | 30.0 | 30.0 | 20.0 |
| NTnC | **62.5** | 40.0 | 37.5 | 50.0 |
| S100A1 | **57.5** | 35.0 | 35.0 | 50.0 |
| S100B | **67.5** | 50.0 | 52.5 | 55.0 |
| STP10 | **70.0** | 65.0 | 67.5 | 60.0 |
| T4L | 67.5 | 77.5 | 67.5 | **80.0** |

### 2.7 Pi-Ensemble structures are consistent with experimental DEER constraints

Finally, we sought to investigate whether Pi-Ensemble can generate structures that span experimentally determined conformational distributions. To this end, we validated Pi-Ensemble predictions against Double Electron-Electron Resonance (DEER) distance distributions using T4 lysozyme as a benchmark system. T4 lysozyme is a bacteriophage-derived enzyme that cleaves bacterial cell wall peptidoglycan to facilitate viral release from infected cells. Owing to its structural stability and conformational adaptability, it has been extensively utilized as a model system for studying protein folding, conformational dynamics, ligand recognition, and protein engineering.^89^ Experimental DEER data were obtained from a study by Islam *et al.* (2013), in which the authors generated 51 DEER distance histograms from spin labels inserted at 34 distinct positions in T4 lysozyme, as shown in Figure 6a.

**Figure 6:**
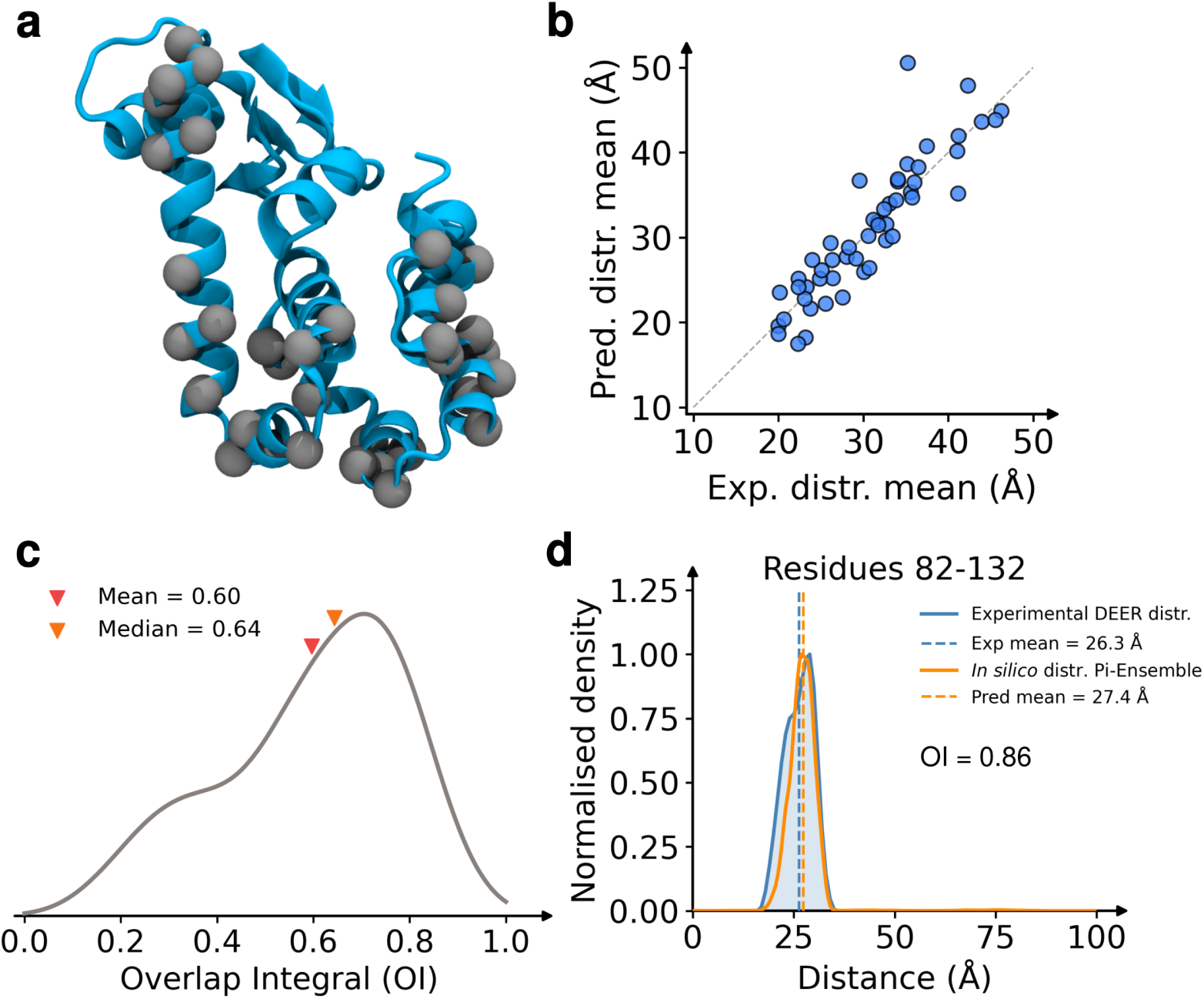
Validation of Pi-Ensemble against Double Electron-Electron Resonance (DEER) spectroscopy data. (a) T4 lysozyme highlighting the 37 spin-labeled sites used as probe locations. (b) Predicted versus experimental mean distances; predicted values are obtained from DEER distance distributions computed from Pi-Ensemble generated structures using DEER-PREdict,^90^ while experimental values are taken from Islam *et al.* (2013).^64^ (c) Kernel density estimate (KDE) of the overlap integral between Pi-Ensemble derived and experimental DEER distributions. (d) Representative example showing the overlap between an experimental DEER distribution and the corresponding Pi-Ensemble prediction for the residue pair 82-132.

To compare Pi-Ensemble predictions against these experimental measurements, we employed DEER-PREdict,^90^ a software framework that predicts DEER distance distributions from ensembles of protein conformations using an established rotamer-library approach. We observe excellent qualitative agreement between the *in silico* predicted distributions derived from Pi-Ensemble structures and the experimentally measured DEER distributions, Figures S13–S16. For a quantitative comparison, we evaluated two complementary metrics after normalizing both distributions. First, we computed the mean distance value of each distribution, which assesses how accurately the Pi-Ensemble predictions reproduce the experimentally observed average distances. Second, we computed the overlap integral (see Methods), which quantifies the similarity between two probability distributions by measuring the shared intersection area between them. An overlap integral value of 1 corresponds to perfect agreement, whereas a value of 0 indicates no overlap between the distributions.

Figure 6b compares the mean values obtained from the experimental DEER distributions for all 51 residue pairs against the corresponding values calculated from Pi-Ensemble-generated structures. We observe excellent agreement between the two sets of values, with a mean absolute error of only 2.32 ÅA. Furthermore, Figure 6c shows the KDE plot of the overlap integral values across all 51 residue pairs. The median overlap integral is 0.64, indicating substantial agreement between the experimental and Pi-Ensemble-derived distance distributions. Finally, Figure 6d highlights a representative residue pair, 82–132, demonstrating a particularly strong overlap between the predicted and experimental distributions.

### 2.8 Pi-Ensemble is implemented as a user-friendly and extensible Python package

Pi-Ensemble is implemented as a modular Python package for generating sequence-guided conformational ensembles between two protein structural anchors. The package couples three interchangeable components: an inverse-folding model that maps a backbone structure to residue-wise amino acid probability distributions, a structure-generation model that maps designed sequences to predicted structures, and an interpolation algorithm that constructs intermediate sequence prompts between anchor states. Each component is exposed through a lightweight registry interface, allowing different pretrained models to be selected from a YAML configuration file without modifying the core workflow.

The current implementation supports ProteinMPNN as the inverse-folding model and ESM3, Boltz, and BioEmu as structure-generation backends. If the two anchors differ in sequence length or contain missing residues, the package aligns the modeled sequences to a user-provided reference sequence, either by computing a pairwise alignment internally or by reading a supplied alignment file. The residue-wise probability distributions are then mapped into this common reference frame before interpolation. Pi-Ensemble provides two complementary interpolation schedules as previously described.

Runs are specified using a YAML file containing the structure-prediction model, inverse-folding model, interpolation algorithm, anchor structures, chain identifiers, reference sequence, and output directory. The package writes predicted structures, model-specific confidence files when available, inverse-folding outputs, and a structured JSON log recording metadata such as interpolation weight, round, source anchors, and edit distance between generated sequences. Optional postprocessing reconstructs the user-specified reference sequence onto generated backbones by retaining backbone coordinates, replacing residue identities, converting to all-atom structures with cg2all, and performing OpenMM energy minimization with selected forcefield parameters. Thus, the software automates the full Pi-Ensemble workflow from two structural anchors to a logged ensemble of predicted intermediate conformations suitable for downstream filtering, visualization, clustering, or MD seeding.

We emphasize that the package is designed for extensibility through abstraction layers that separate model-specific wrappers from the interpolation algorithms or the user interface. As a result, new inverse-folding or structure-prediction models can be incorporated by implementing a lightweight wrapper that conforms to the corresponding abstract interface and registering the model under a user-defined name. Once registered, the model can be selected directly from the YAML configuration file without modifying the core Pi-Ensemble workflow. These abstract base classes define the expected inputs and outputs for each model type, ensuring consistency across contributed wrappers while allowing the framework to remain compatible with future advances in protein sequence design and structure generation. More details are available in the code repository (see Data availability).

## 3 Discussion

In this work, we introduced Pi-Ensemble, a sequence-guided framework for generating interpolated protein conformational ensembles between structural templates. Rather than interpolating directly in Cartesian coordinate space or retraining a generative model on MD trajectories, Pi-Ensemble couples two pretrained model classes with orthogonal objectives: an inverse-folding model that proposes residue identity distributions compatible with a given backbone, and a structure-prediction model that maps the resulting sequence prompts back to three-dimensional structures. By interpolating between inverse-folding probability distributions and iteratively feeding predicted structures back into the design cycle, Pi-Ensemble generates conformations that bridge input anchor states without requiring system-specific training. Across the systems examined here, this strategy produced structurally plausible intermediate conformations, recovered transition pathways observed in large-scale MD simulations, provided effective seed structures for accelerated conformational sampling, generalized across diverse protein families, and yielded ensembles consistent with experimental DEER distance distributions.

The interaction between inverse folding and structure prediction provides a useful way to interpret why Pi-Ensemble works when it succeeds. The inverse-folding model introduces backbone-conditioned biochemical information: given a backbone pose, it proposes residue identities that are compatible with, or may stabilize, that pose. The structure-prediction model then acts as a sequence–structure evaluator: given the designed sequence, it identifies a backbone conformation that is plausible under the learned relationships between sequence and structure. Because these models are trained with different objectives, their coupling creates a form of bootstrapping. Importantly, this process does not require the structure-prediction model to have been explicitly trained to generate transition paths or Boltzmann-weighted ensembles. Instead, conformational diversity emerges from the interplay between two models trained on related structural data but optimized for different tasks.

A central observation motivating this work is that pretrained structure-prediction models can be steered toward conformationally diverse outputs by modifying the sequence prompts used to query them. Previous approaches have exploited the sensitivity of MSA-based structure-prediction models to input perturbations, primarily MSA subsampling, to generate alternative conformations. However, structure-prediction models are generally trained to penalize sensitivity to different MSAs corresponding to the same sequence.^19^ Pi-Ensemble follows a conceptually related but mechanistically distinct strategy. Instead of perturbing the MSA, our method exploits the numerical sensitivity of inverse-folding models. In particular, ProteinMPNN, or related methods, can assign different residue preferences to backbone conformations that may correspond to the same natural protein sequence. These differences encode local and nonlocal sequence features compatible with each backbone state. By interpolating between such residue probability distributions, Pi-Ensemble constructs sequence prompts that are not necessarily natural sequences, but that retain structural information about the two endpoint conformations. These prompts can then bias structure-prediction models toward conformations that span the region between the anchors. More broadly, our results suggest that the numerical sensitivity of inverse-folding models to structural perturbations can be a useful feature rather than merely a source of instability, because it encodes backbone-dependent sequence preferences that can be exploited to guide conformational exploration. This perspective may inform future strategies for structure-based design and sequence-guided conformational sampling.

Pi-Ensemble should not be interpreted as a replacement for physics-based simulation or experimental ensemble determination. The generated structures are not guaranteed to follow a physical transition pathway, and the density of generated conformations should not be interpreted directly as a thermodynamic probability distribution. The interpolation coordinate *λ* operates in inverse-folding probability space rather than along a physical reaction coordinate. Consequently, while Pi-Ensemble can generate plausible intermediates and useful starting points for simulation, additional filtering, minimization, MD relaxation, or comparison with experimental observables may be required before drawing quantitative conclusions about free energies, state populations, or kinetics. In this sense, Pi-Ensemble is best viewed as a conformational proposal mechanism: it suggests structurally plausible regions of conformational space that can then be refined or reweighted using physics-based or experimentally guided approaches. In particular, our method may be used in combination with physically-motivated approaches such as the string method^91^ or energy-guided techniques such as DeepPath.^37^

The method also inherits limitations from its constituent models. Inverse-folding models such as ProteinMPNN can produce residue distributions that are useful for backbone-conditioned design, but they may also favor pathological or non-natural sequence features in some regimes, including low-complexity segments, poly-alanine stretches, or KE-rich regions.^92^ When such sequences are passed to a structure-prediction model, they can induce artifacts such as disordered regions or artificial helices. This issue is particularly important because Pi-Ensemble intentionally moves away from the native sequence in order to probe alternative conformations. Although the native sequence is reconstructed onto the final generated backbones before downstream use, the intermediate sequence prompts still determine which backbone structures are proposed. Thus, failures in the sequence-prompting stage can propagate into the structural ensemble. We observed related behavior when attempting to use BioEmu as a structure-generation backend in Pi-Ensemble. Although BioEmu is explicitly trained to produce conformational ensembles, many structures generated from interpolated sequence prompts were filtered because of atomic clashes. This suggests that not all structure-generation models are equally compatible with sequence-space interpolation, and that models trained for ensemble generation still require physically informed steering or filtering when queried with artificial interpolated sequences.

Comparisons across structure-generation backends further suggest that different models capture distinct modes of conformational diversity (Figures S8–S12). Across the systems analyzed here, ESM3 appeared to recover greater interpolated diversity for smaller proteins whose transitions involve intradomain secondary-structure rearrangements, such as NTnC and S100B. In contrast, ESMFold2 more effectively sampled interdomain hinge-bending motions in systems such as LBP and ALBP. BioEmu outputs tended to resemble those obtained with Pi-Ensemble coupled to ESMFold2, whereas Pi-Ensemble coupled to ESM3 sometimes revealed intermediates that were less accessible to the other backends. These observations suggest that model choice can strongly influence the regions of conformational space explored by sequence-guided interpolation, and that Pi-Ensemble may provide a useful alternative when AlphaFold2-derived or ensemble-generation models fail to capture the intended conformational change.

Moreover, the limitations observed in some of the tested examples help clarify both the current domain of applicability of Pi-Ensemble and the most important directions for future development. The method is best suited to proteins for which two or more known, predicted, or homologous structural templates define a meaningful conformational change, particularly when the goal is to generate plausible intermediates for visualization, hypothesis generation, adaptive or enhanced sampling (e.g., umbrella sampling or string method initialization), or other forms of distributed MD seeding. It may be less reliable when the anchors are poorly resolved, when the transition involves extensive unfolding, ligand-dependent stabilization, changes in oligomeric state, membrane or cofactor effects not represented by the structure-prediction model, or when one endpoint lies outside the learned structural distribution of the underlying models. In these cases, interpolated sequence prompts may fail to encode a coherent structural path, causing generated structures to collapse toward one anchor or drift into nonphysical regions of conformational space. A practical strength of the framework is that these limitations can be addressed modularly: although the present implementation uses ProteinMPNN with ESM3, Boltz, or ESMFold2, Pi-Ensemble separates inverse folding, interpolation scheduling, structure generation, and postprocessing, allowing improved models to be incorporated as they become available. Future work should therefore focus on regularizing interpolated sequence prompts, penalizing low-complexity or pathological designs, adding physics-informed steering or filtering during generation, and coupling generated structures with MD, MSMs, experimental restraints, or learned reweighting schemes to assign more quantitative ensemble weights. Together, these developments could extend Pi-Ensemble from a conformational proposal method into a more complete framework for estimating protein conformational landscapes. Overall, the results presented here demonstrate that sequence-guided interpolation between structural templates is a promising strategy for accelerating investigations of protein dynamics, especially when long-timescale MD simulations or task-specific model retraining are impractical.

## 4 Methods

### 4.1 Performance metrics

#### 4.1.1 Root Mean Square Deviation

Unless otherwise specified, the root mean square deviation (RMSD) was calculated for C*_α_*atoms following structural alignment according to

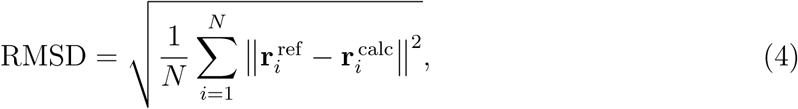

where *N* is the number of C*_α_*atoms, 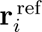 denotes the position vector of atom *i* in the reference structure, and **r** ^calc^ denotes the position vector of the corresponding atom in the aligned structure.

#### 4.1.2 Relative Entropy

Relative entropy between two transition probability matrices derived from MSMs was computed to quantify convergence. Specifically, we employed the relative entropy metric introduced by Bowman *et al.*,^93^ defined as

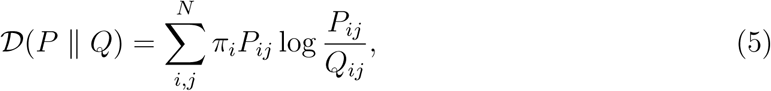

where *P* and *Q* denote the transition probability matrices of the reference and test MSMs, respectively, and *π_i_* represents the stationary (equilibrium) probability of state *i*. Lower values of *D* indicate greater similarity between the test and reference transition matrices. The reference MSM corresponds to the converged ground-truth MSM, while the test MSM corresponds to the MSM constructed at increasing sampling time. To ensure consistency between *P* and *Q*, the state definitions from the ground-truth MSM were imposed on all test MSMs, thereby preserving the ordering and dimensionality of the transition matrices. Additionally, a uniform prior of 1/*N*, where *N* is the number of states, was incorporated into the test MSM count matrix as pseudocounts prior to row normalization. This prior serves two purposes. First, it prevents Eq. 5 from becoming undefined in cases where zero transition probabilities are encountered. Second, before sufficient sampling is observed, transitions from a given state are assumed to be uniformly distributed across all possible target states, motivating the choice of a uniform prior.

#### 4.1.3 Overlap Integral

Agreement between experimental DEER distance distributions and Pi-Ensemble predicted distributions from DEERpredict was quantified using the overlap integral (OI). For a given spin-label pair, the OI is defined as

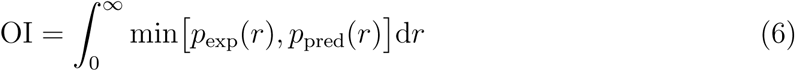

where *p*_exp_(*r*) is the normalized experimental distance probability distribution and *p*_pred_(*r*) is the ensemble-mean distribution calculated by averaging the individual probability distributions within the predicted ensemble. Prior to evaluation, distributions were interpolated onto a common distance grid and renormalized numerically as *p*(*r*) = *p*(*r*)/ ∫ *p*(*r*) d*r*. The OI ranges from 0 (no overlap) to 1 (perfect agreement). All integrals were evaluated numerically using the trapezoidal rule.

### 4.2 Datasets

The MD dataset for the AtSWEET13 system was obtained from Weigle *et al.* (2024).^15^ Only the *apo* trajectories, comprising a total of 132 *µ*s of simulation data, were used in this study. The gating distance features used in Figure 2 were also derived from this dataset. The MD dataset for the GlnBP system was sourced from the Open Research Data Repository of the Max Planck Society (Edmond). Specifically, we used the dataset Weikl, Thomas (2026), Atomistic MD trajectories of the glutamine-binding protein GlnBP, version 1.^62^ The benchmark set of 18 proteins shown in Figure 5 was adapted from Li *et al.* (2023).^63^

### 4.3 Molecular Dynamics

Pi-Ensemble-generated structures were clustered using the Regular Space clustering algorithm implemented in the Deeptime library,^94^ with a minimum distance parameter (dmin) of 0.1.

All MD simulations were performed using OpenMM 8.2.0.^60^ Systems were first energy minimized, followed by a 1 ns equilibration in the NVT ensemble. During this stage, the temperature was gradually increased from 5 K to 300 K. Protein backbone atoms were restrained using a harmonic force constant of 1.0 kcal/mol ÅA*^−^*^2^ throughout the NVT equilibration.

Subsequently, systems were equilibrated for 1 ns in the NPT ensemble, consisting of 0.5 ns with backbone restraints followed by 0.5 ns without restraints. Long-range electrostatic interactions were treated using the particle mesh Ewald (PME) method with a nonbonded cutoff of 1.2 nm. Hydrogen-containing bonds were constrained, and water molecules were maintained as rigid bodies. Simulations employed a 2 fs integration timestep and a Langevin thermostat with a friction coefficient of 2.8284 ps*^−^*^1^. Pressure was maintained at 1 atm during NPT simulations.

Trajectory frames were saved every 100 ps for downstream analysis. Production simulations used for the final analyses were also conducted in the NPT ensemble using the same simulation parameters. Simulation durations for both AdK and GlnBP systems are described in Section 2.5 (Results). The CHARMM36m force field^95^ was used for proteins, while the TIP3P water model^96^ was used for solvent molecules.

### 4.4 Markov State Models

Two collective variables describing the conformational state of AdK were used as features for MSM construction: the NMP-domain angle (*θ*_NMP_) and the LID-domain angle (*θ*_LID_). Each angle was calculated using the centers of geometry of backbone and C*β* atoms across three domain-defining residue groups. Specifically, *θ*_NMP_ was defined using residues 114–124, 89–99, and 34–54, whereas *θ*_LID_ was defined using residues 178–184, 114–124, and 124–152. Featurization was performed using MDTraj.^97^

The resulting two-dimensional angle features were used as input for MSM construction. Trajectories were discretized into *k* = 50 microstates using mini-batch *k*-means clustering with *k*-means++ initialization. Transition count matrices were estimated using a sliding-window approach, and implied timescales were calculated over lag times ranging from 1 to 49 steps (0.1–4.9 ns) to evaluate Markovian behavior. The final MSMs was constructed at a lag time of *τ* = 30 steps (3 ns) using maximum-likelihood estimation on the largest connected submodel. Model validity was further assessed using Chapman-Kolmogorov (CK) tests with *n* = 3 metastable states. All MSM analyses were performed using Deeptime.^94^ Implied timescale plots for Pi-Ensemble seeded simulations and anchors-only seeded simulations are shown in Figure S4 and Figure S6 respectively. CK test plots for Pi-Ensemble seeded simulations and anchors-only seeded simulations are shown in Figure S5 and Figure S7.

### 4.5 Run settings

Pi-Ensemble with serial interpolation was run for 20 rounds with *λ ∈* [0, 1]. The batched interpolation variant was also run for 20 rounds, but we note that the number of output structures is input-dependent for this algorithm; it depends on the number of unique generated sequences or critical *λ* values. BioEmu was executed with filter samples=True to remove physically invalid structures, including those containing steric clashes. ProteinMPNN was used for sequence prediction with a low-temperature setting (*T* = 0.1), a fixed seed of 1, batch size of 1. For structure prediction, ESM3 was run on a GPU device with temperature 0.7 and 8 sampling steps, while Boltz was executed on GPU with one recycling step, a single diffusion sample, one preprocessing thread, and MSA retrieval in server mode. Sample configuration files for all methods are available in the accompanying GitHub repository.

## Data availability

Data used to produce the figures in this study are available in the Illinois Data Bank at https://doi.org/10.13012/B2IDB-6902835_V1 and the corresponding references.

Code required to reproduce the results is available at https://github.com/ShuklaGroup/ Pi-Ensemble.

## Acknowledgement

This work was supported by the Army Research Office under Cooperative Agreement Number W911NF-22-2-0246. Authors acknowledge support from the National Institutes of Health (Award No. R35GM142745). This material is also based upon work supported by the National Science Foundation under Award No. 2435360.

The views and conclusions contained in this document are those of the authors and should not be interpreted as representing the official policies, either expressed or implied, of the Army Research Office or the U.S. Government. The U.S. Government is authorized to reproduce and distribute reprints for Government purposes notwithstanding any copyright notation herein.

We thank Dr. Austin Weigle for providing MD data for the AtSWEET13 system.

The authors acknowledge the use of GPT-5.5 (OpenAI) to assist with improving the language, flow, and clarity of the manuscript. The authors reviewed, edited, and approved all AI-assisted text and take full responsibility for the content of this publication. The authors also used Codex (OpenAI) as an AI-assisted coding tool during software development and code production.

## Supporting information

Figures S1–S16.

## Author contributions

H.N. and D.E.K. contributed equally to this work. H.N., D.E.K., and D.S. designed the study. H.N., D.E.K., and Y.Z. developed the codebase. H.N., D.E.K., and Y.Z. conducted the experiments and performed the analyses. H.N., D.E.K., and Y.Z. wrote the manuscript with input from A.D.B.L. and D.S. A.D.B.L. and D.S. secured funding for the project.

