## Supplementary Information for "Pi-Ensemble: Sequence-guided generation of interpolated protein conformational ensembles"

### 1 Supporting Tables

Table S1: Source structures for 18 test cases presented in Figure 5.

| # | Protein | Function | Size | Conformational change | Anchor 1 | Anchor 2 | RMSD (Å) | Missing residues |
|---|---|---|---|---|---|---|---|---|
| 1 | 48G7 | Immunoglobin 48G7 germline | 217 | Domain rotation | 1AJ7 | 2RCS | 7.21 | No |
| 2 | AK | Adenylate kinase | 214 | Combination | 2ECK | 4AKE | 6.95 | No |
| 3 | AP4A | Ap4A hydrolase | 165 | Single-domain helices and loops | 1F3Y | 1JKN | 3.68 | No |
| 4 | ASCT2 | Neutral amino acid transporter | 447 | Transmembrane helices and loops | 6GCT | 7BCQ | 11.40 | Yes |
| 5 | CNBD | Nucleotide-activated $K^+$ channel binding domain | 142 | Single-domain helices and loops | 2K0G | 2KXL | 5.16 | No |
| 6 | DAHPS | DAHP synthase | 343 | Domain rotation | 1RZM | 1VR6 | 10.28 | Yes |
| 7 | GlnBP | Glutamine-binding protein | 223 | Hinge bending | 1GGG | 1WDN | 5.40 | Yes |
| 8 | GroEL | 60 kDa chaperonin | 525 | Combination | 1AON | 4HEL | 12.78 | Yes |
| 9 | HSPD1 | Human mitochondrial chaperonin | 527 | Combination | 4PJ1 | 7AZP | 13.97 | Yes |
| 10 | LBP | Leucine-binding protein | 346 | Hinge bending | 1USG | 1USI | 7.14 | No |
| 11 | ALBP | D-allose-binding protein | 288 | Hinge bending | 1GUD | 1RPJ | 4.45 | No |
| 12 | Mhp1 | Hydantoin transport protein | 466 | Transmembrane helices and loops | 2JLN | 2X79 | 3.52 | Yes |
| 13 | NTnC | Regulatory domain of troponin C | 90 | Single-domain helices and loops | 1SKT | 1TNQ | 6.03 | No |
| 14 | NTRC | Receiver domain of transcriptional enhancer-binding protein | 124 | Single-domain helices and loops | 1KRX | 1NTR | 3.70 | No |
| 15 | S100A1 | S100 metal-binding protein | 93 | Single-domain helices and loops | 1K2H | 1ZFS | 6.10 | No |
| 16 | S100B | S100 calcium-binding protein | 92 | Single-domain helices and loops | 1SYM | 1XYD | 6.06 | No |
| 17 | STP10 | Sugar transport protein 10 | 492 | Transmembrane helices and loops | 7AAQ | 7AAR | 5.33 | Yes |
| 18 | T4L | Lysozyme | 164 | Hinge bending | 172L | 2LZM | 3.32 | No |

#### 2 Supporting Figures

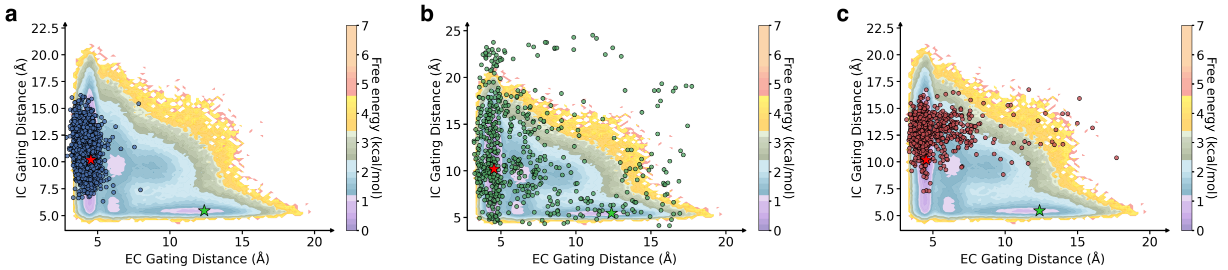


Figure S1: Projection of (a) BioEmu (b) Pi-Ensemble (Boltz), serial interpolation (c) Pi-Ensemble (ESM3) structures onto the two-dimensional space defined by intra- and extracellular gating distances for AtSWEET13.

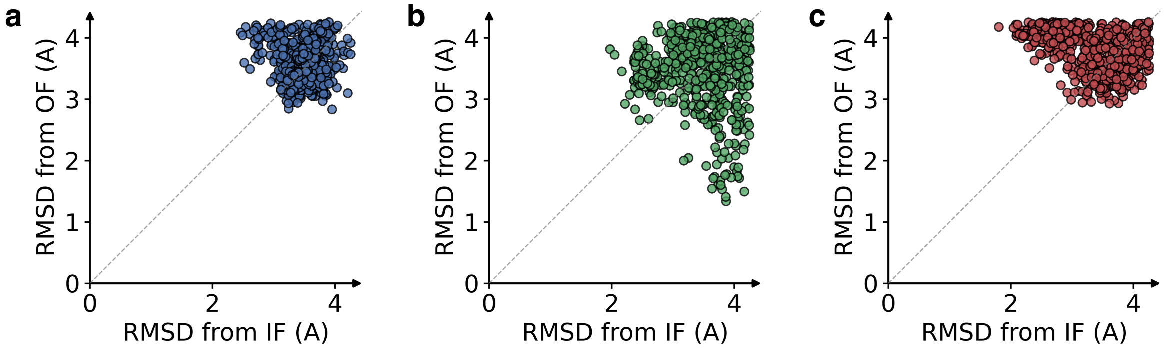


Figure S2: Scatter plot showing projection of (a) BioEmu (b) Pi-Ensemble (Boltz) (c) Pi-Ensemble (ESM3) structures onto the two-dimensional RMSD space, where the axes show distance from each anchor structure respectively, for AtSWEET13.

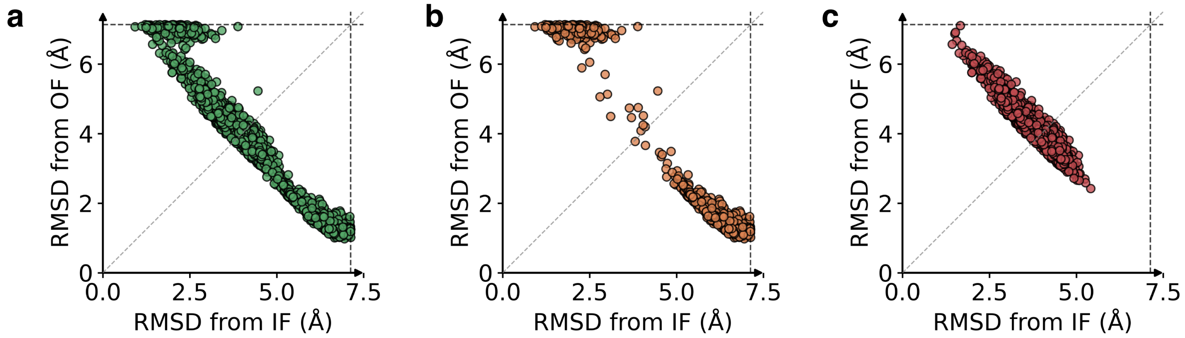


Figure S3: Scatter plots showing projection of structures generated by Pi-Ensemble for AdK system. (a) Combined structures from Pi-Ensemble (Boltz) and Pi-Ensemble (ESM3), used for seeding Pi-Ensemble simulations. (b) Structures from Pi-Ensemble (Boltz). (c) Structures from Pi-Ensemble (ESM3).

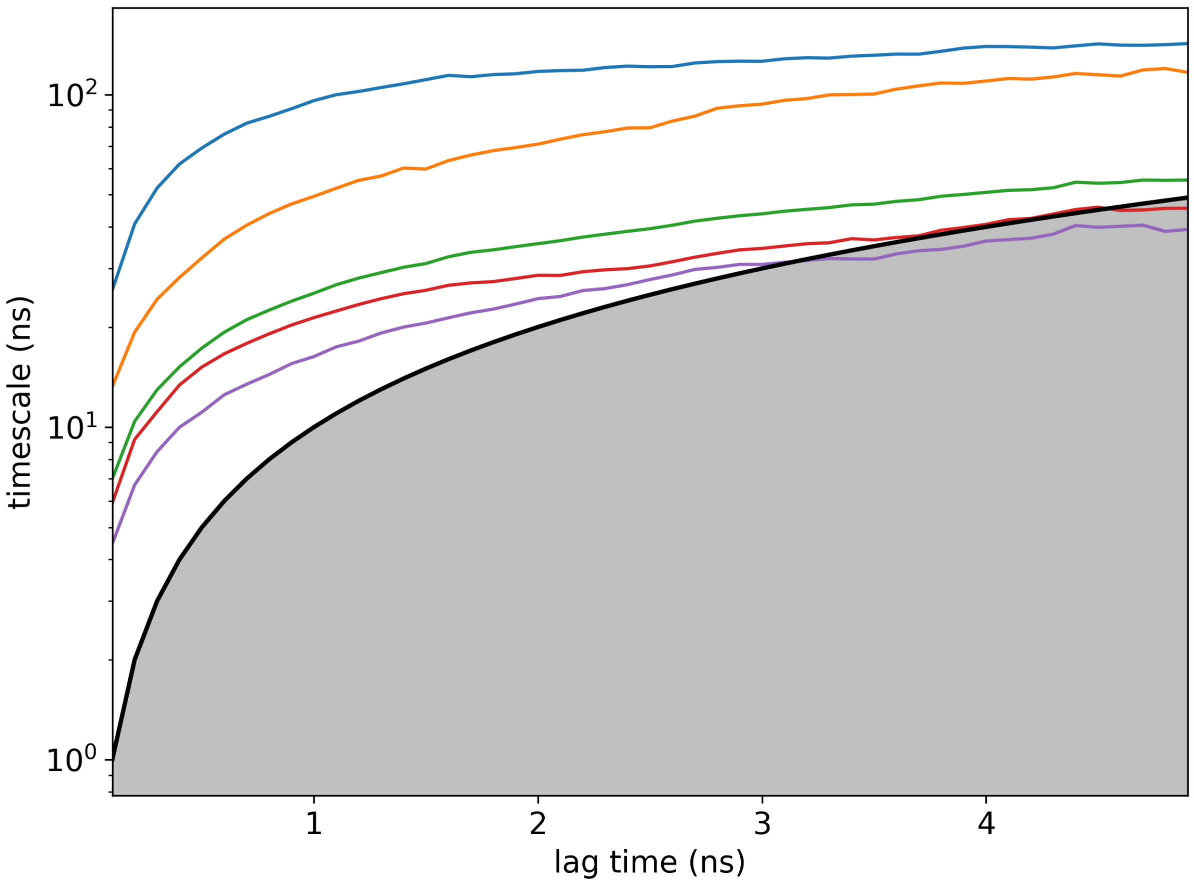


Figure S4: Implied timescales plot for Pi-Ensemble seeded simulations for AdK system.

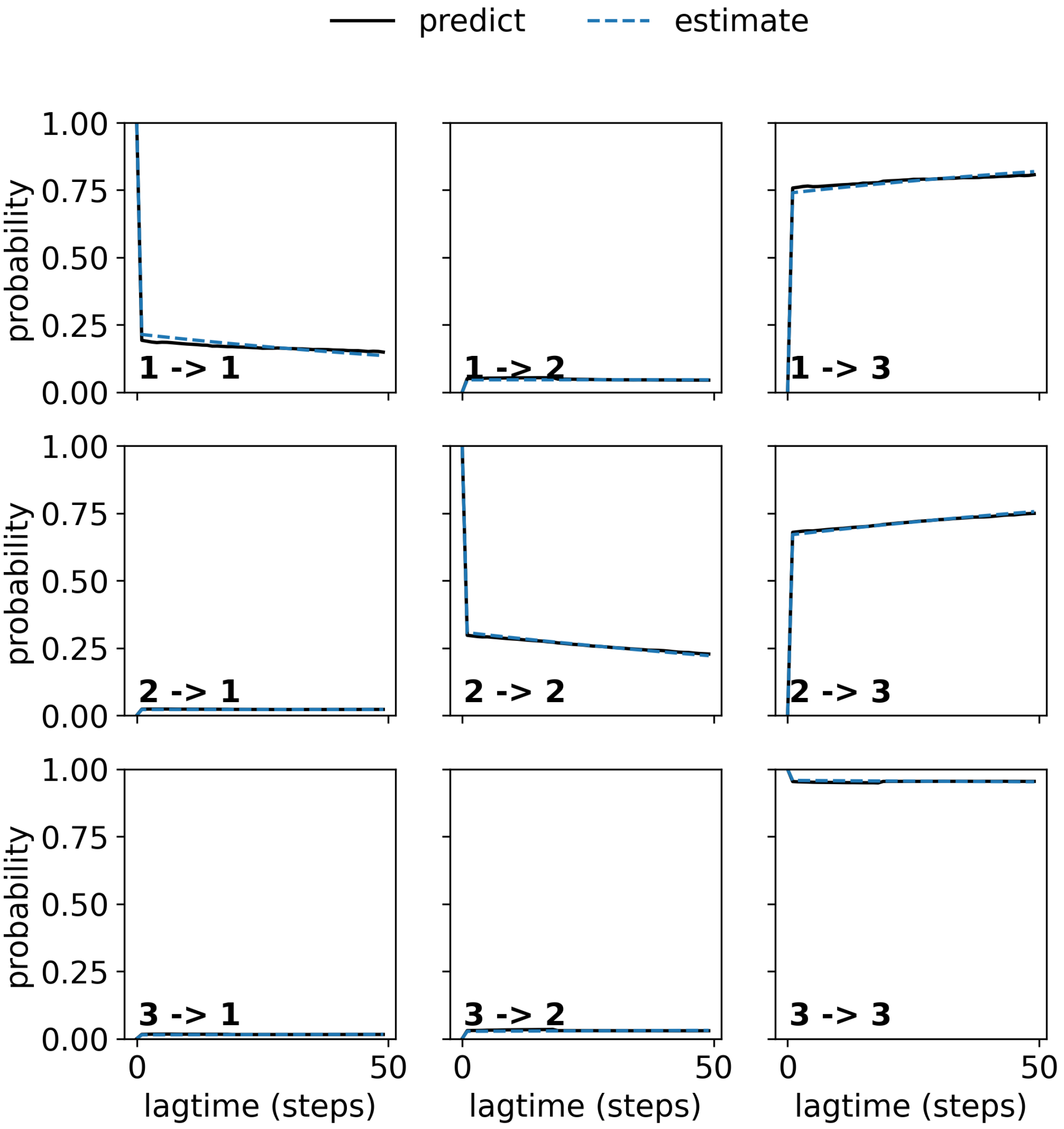


Figure S5: Chapman-Kolmogorov test plot for Pi-Ensemble seeded simulations for AdK system.

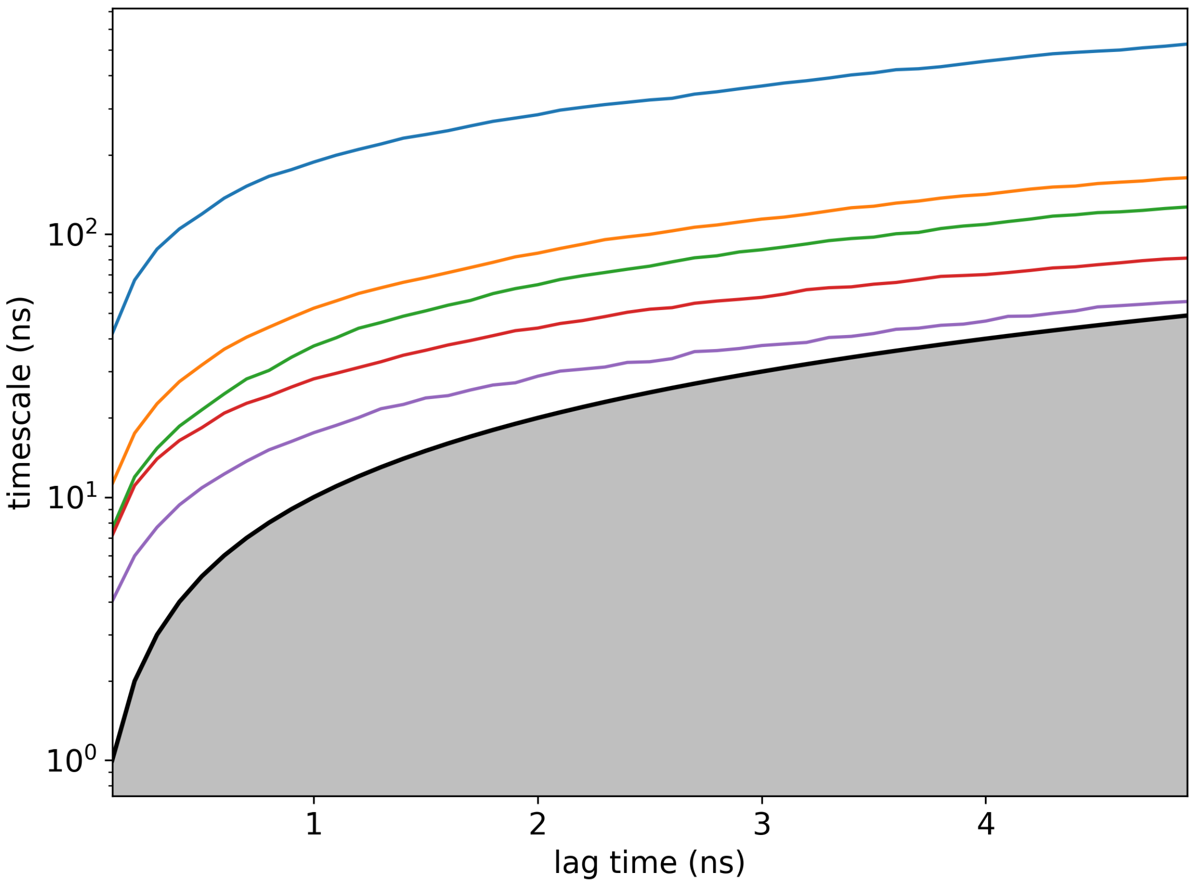


Figure S6: Implied timescales plot for simulations seeded from anchors (PDB structure templates) for AdK system.

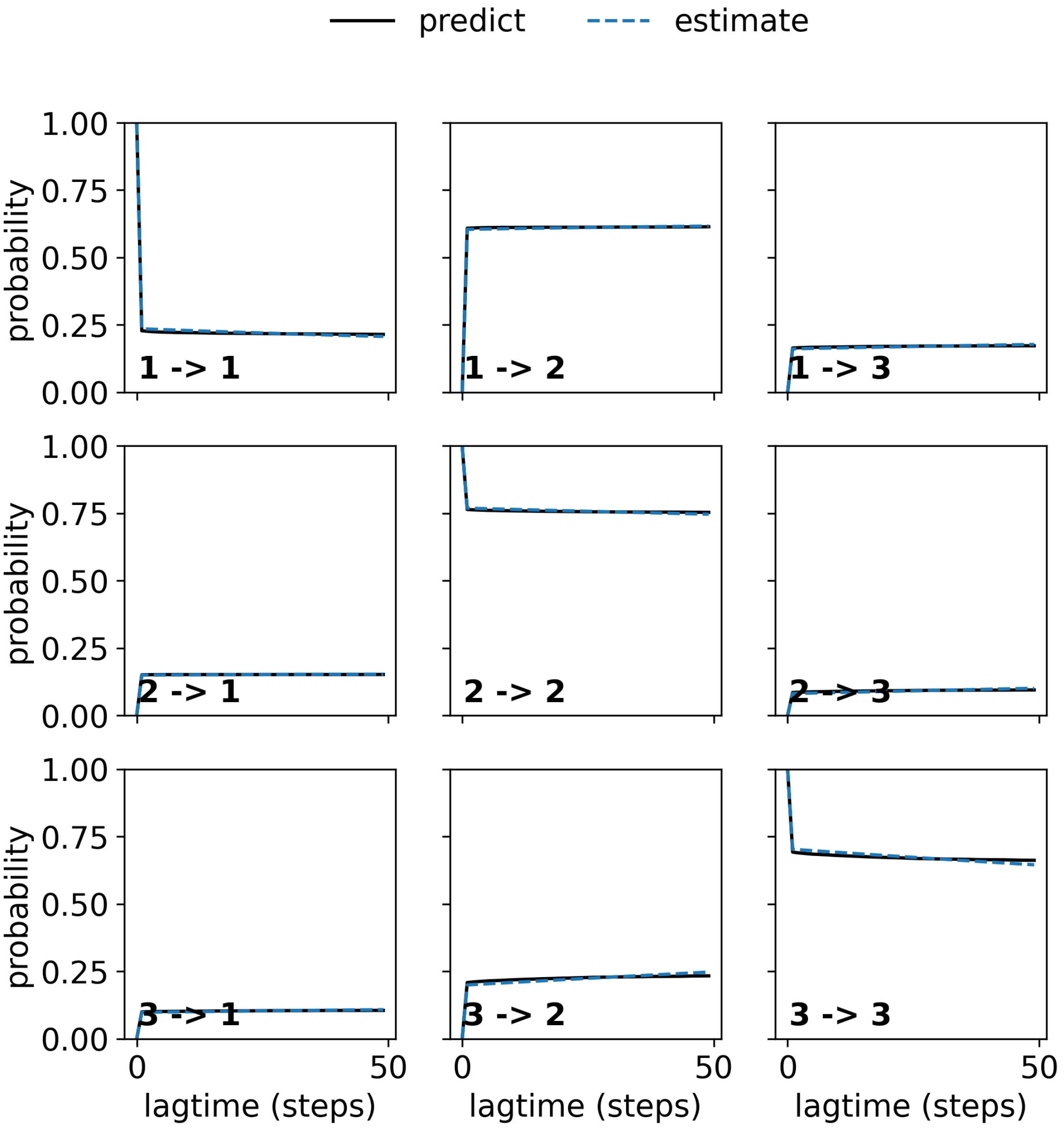


Figure S7: Chapman-Kolmogorov test plot for simulations seeded from anchors (PDB structure templates) for AdK system.

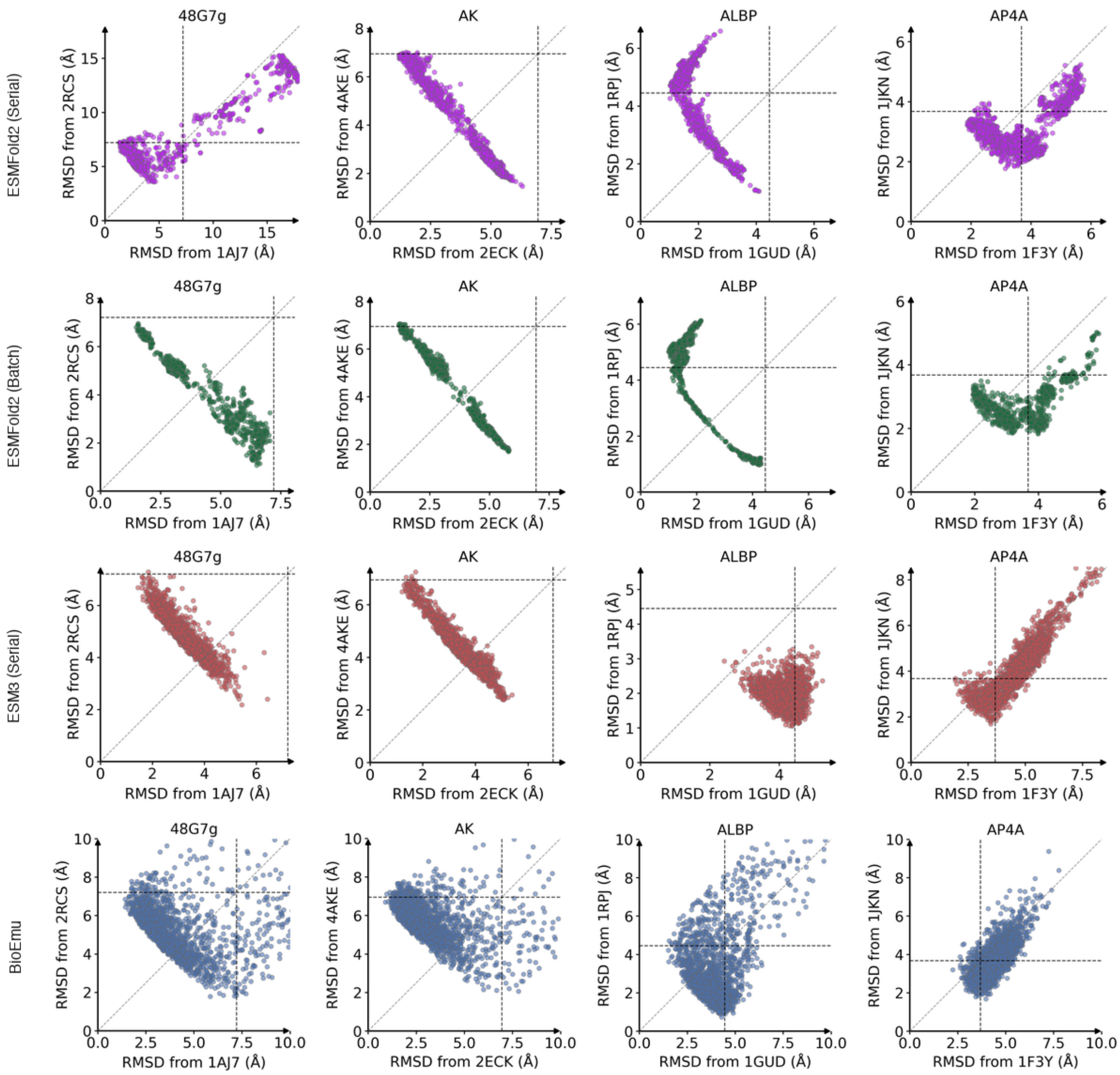


Figure S8: Comparison of Pi-Ensemble structures for a benchmark set of 18 proteins using different structure prediction models and $\lambda$ scheduling strategies. ESM3 (Serial) is shown in red, ESMFold2 (Batch) in purple, ESMFold2 (Serial) in green, and BioEmu (blue) is included as a reference. Results for proteins 1–4 of the 18-protein benchmark set.

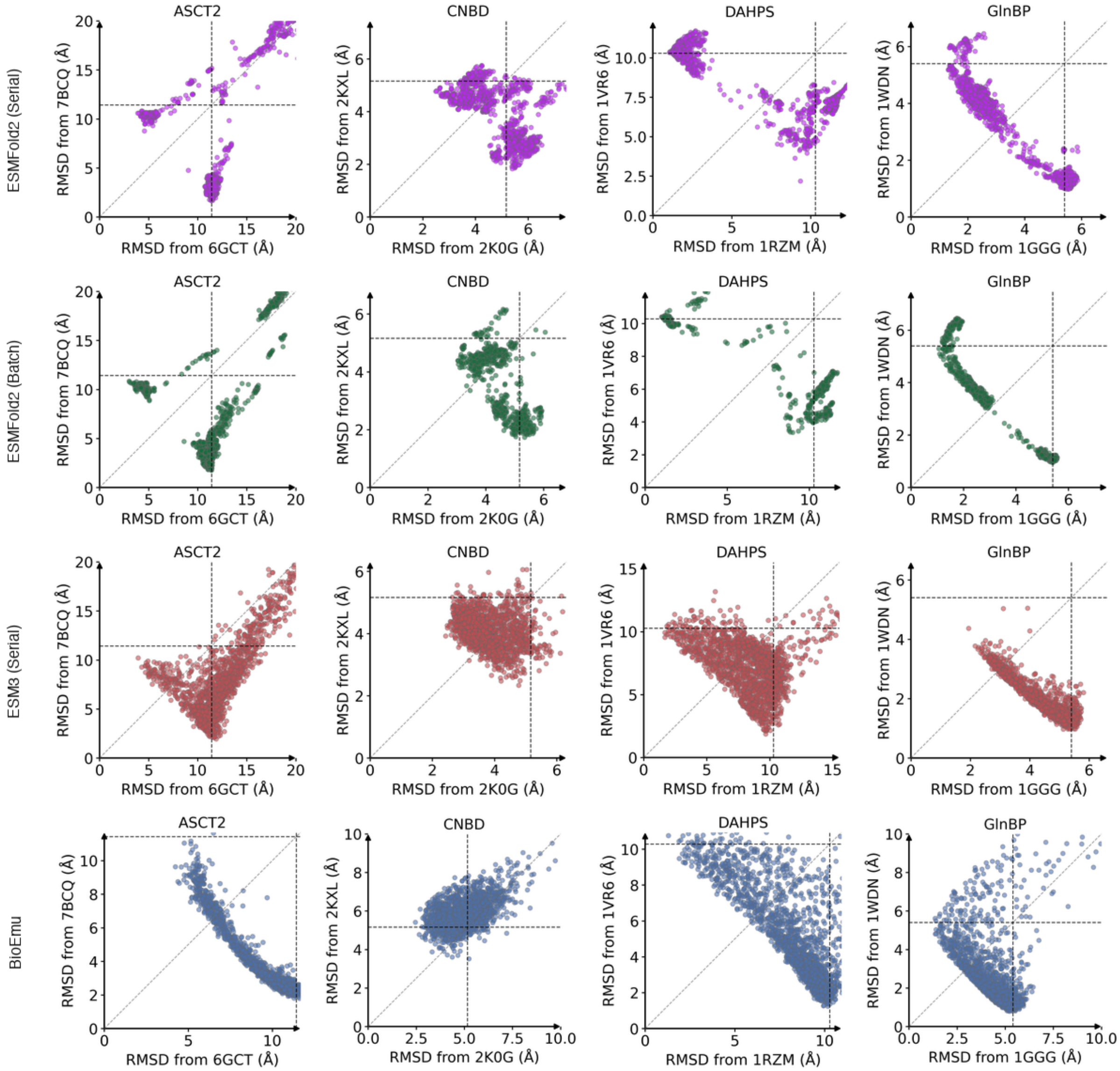


Figure S9: Comparison of Pi-Ensemble structures for a benchmark set of 18 proteins using different structure prediction models and $\lambda$ scheduling strategies. ESM3 (Serial) is shown in red, ESMFold2 (Batch) in purple, ESMFold2 (Serial) in green, and BioEmu (blue) is included as a reference. Results for proteins 5–8 of the 18-protein benchmark set.

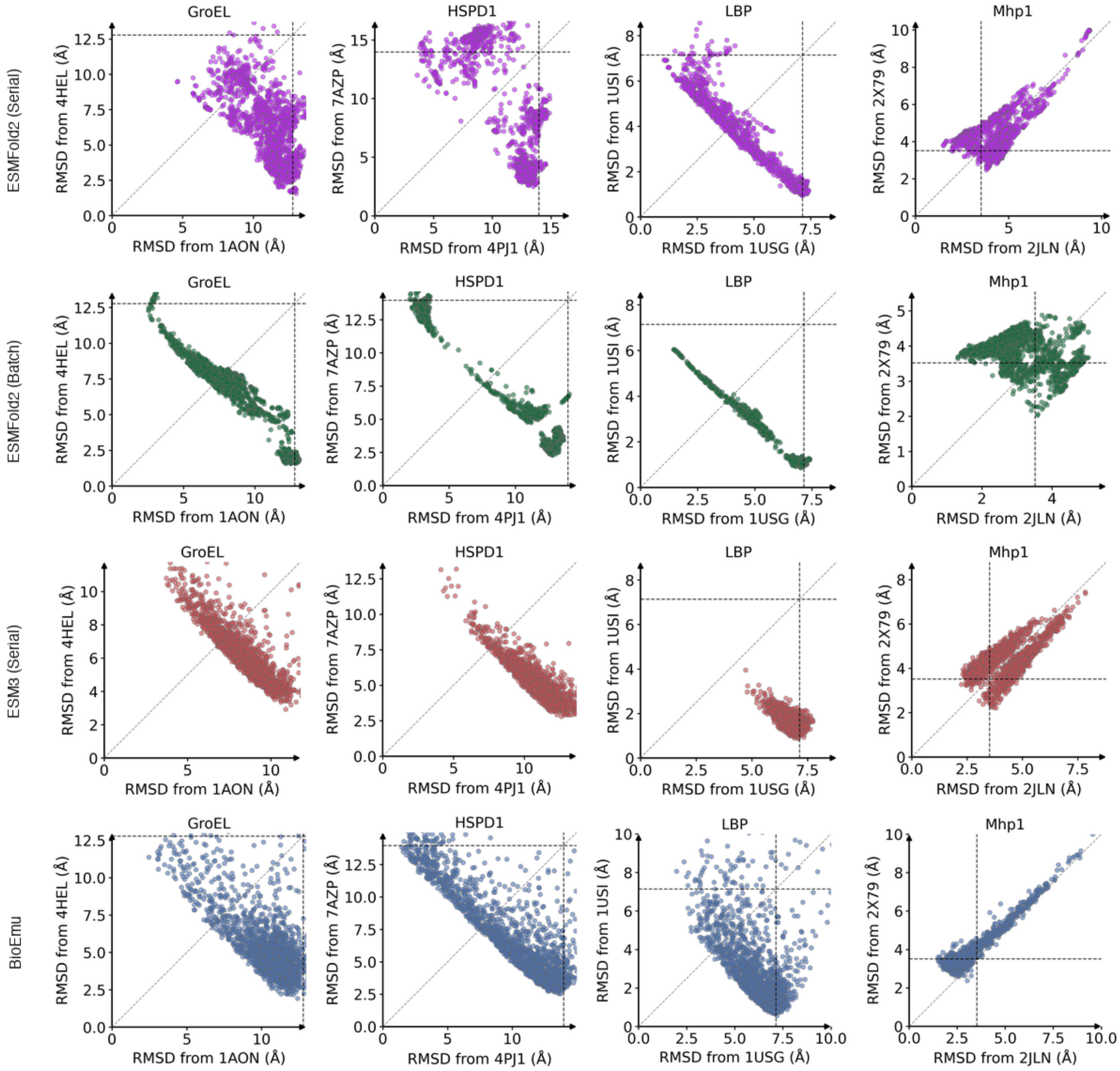


Figure S10: Comparison of Pi-Ensemble structures for a benchmark set of 18 proteins using different structure prediction models and $\lambda$ scheduling strategies. ESM3 (Serial) is shown in red, ESMFold2 (Batch) in purple, ESMFold2 (Serial) in green, and BioEmu (blue) is included as a reference. Results for proteins 9–12 of the 18-protein benchmark set.

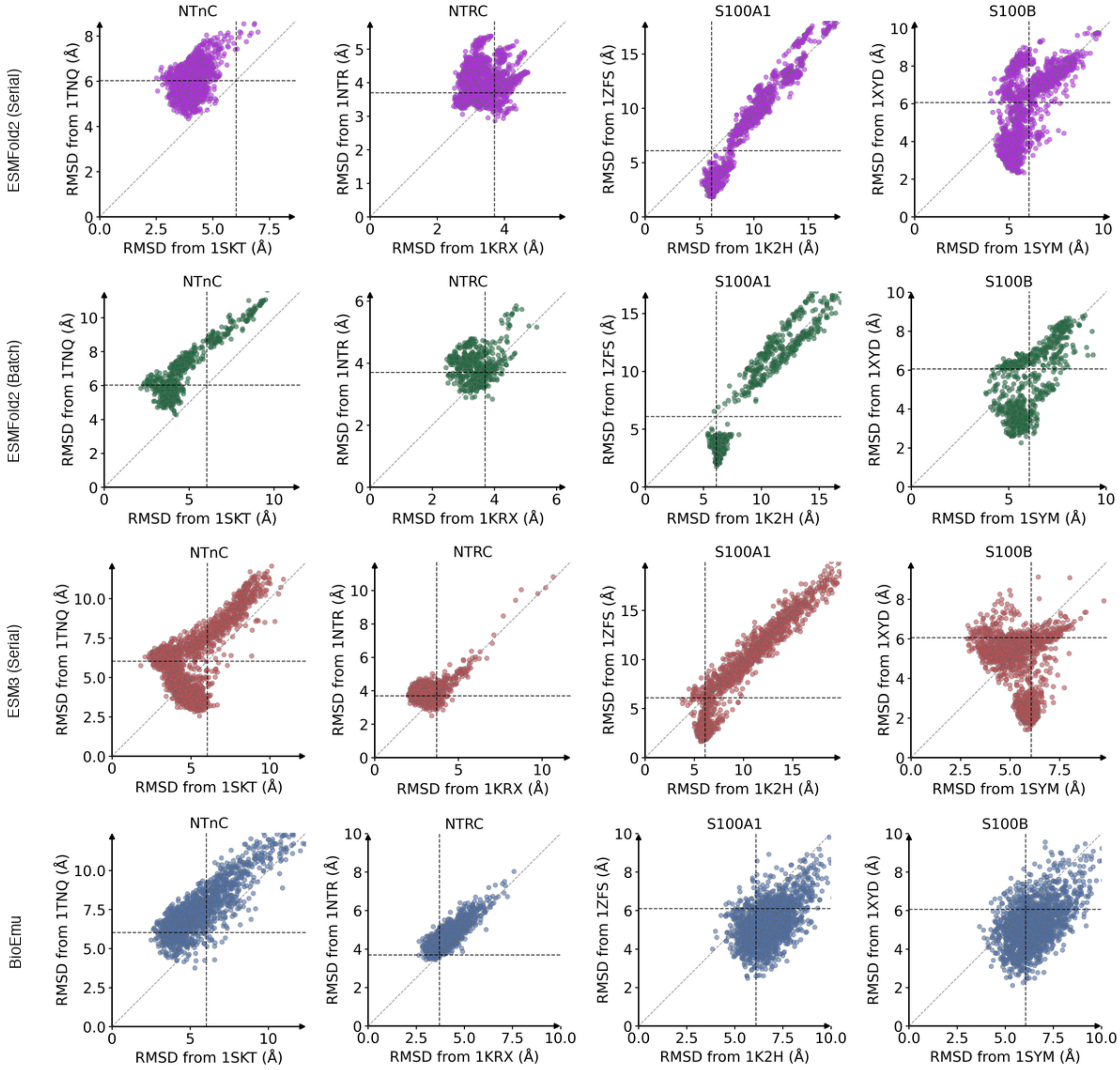


Figure S11: Comparison of Pi-Ensemble structures for a benchmark set of 18 proteins using different structure prediction models and $\lambda$ scheduling strategies. ESM3 (Serial) is shown in red, ESMFold2 (Batch) in purple, ESMFold2 (Serial) in green, and BioEmu (blue) is included as a reference. Results for proteins 13–16 of the 18-protein benchmark set.

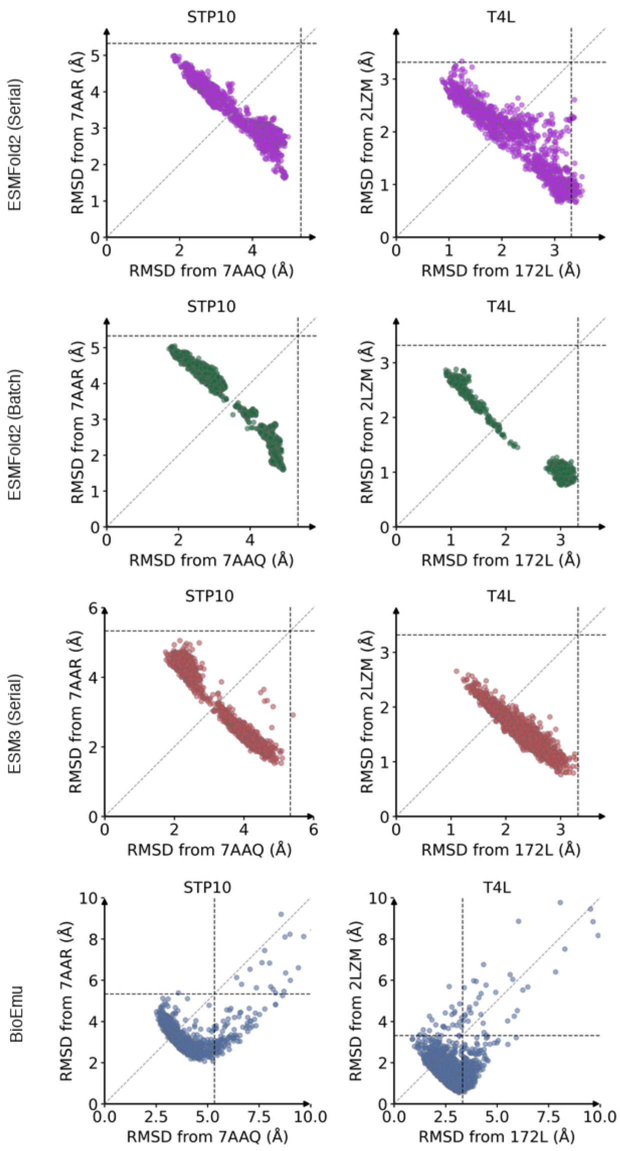


Figure S12: Comparison of Pi-Ensemble structures for a benchmark set of 18 proteins using different structure prediction models and $\lambda$ scheduling strategies. ESM3 (Serial) is shown in red, ESMFold2 (Batch) in purple, ESMFold2 (Serial) in green, and BioEmu (blue) is included as a reference. Results for proteins 17–18 of the 18-protein benchmark set.

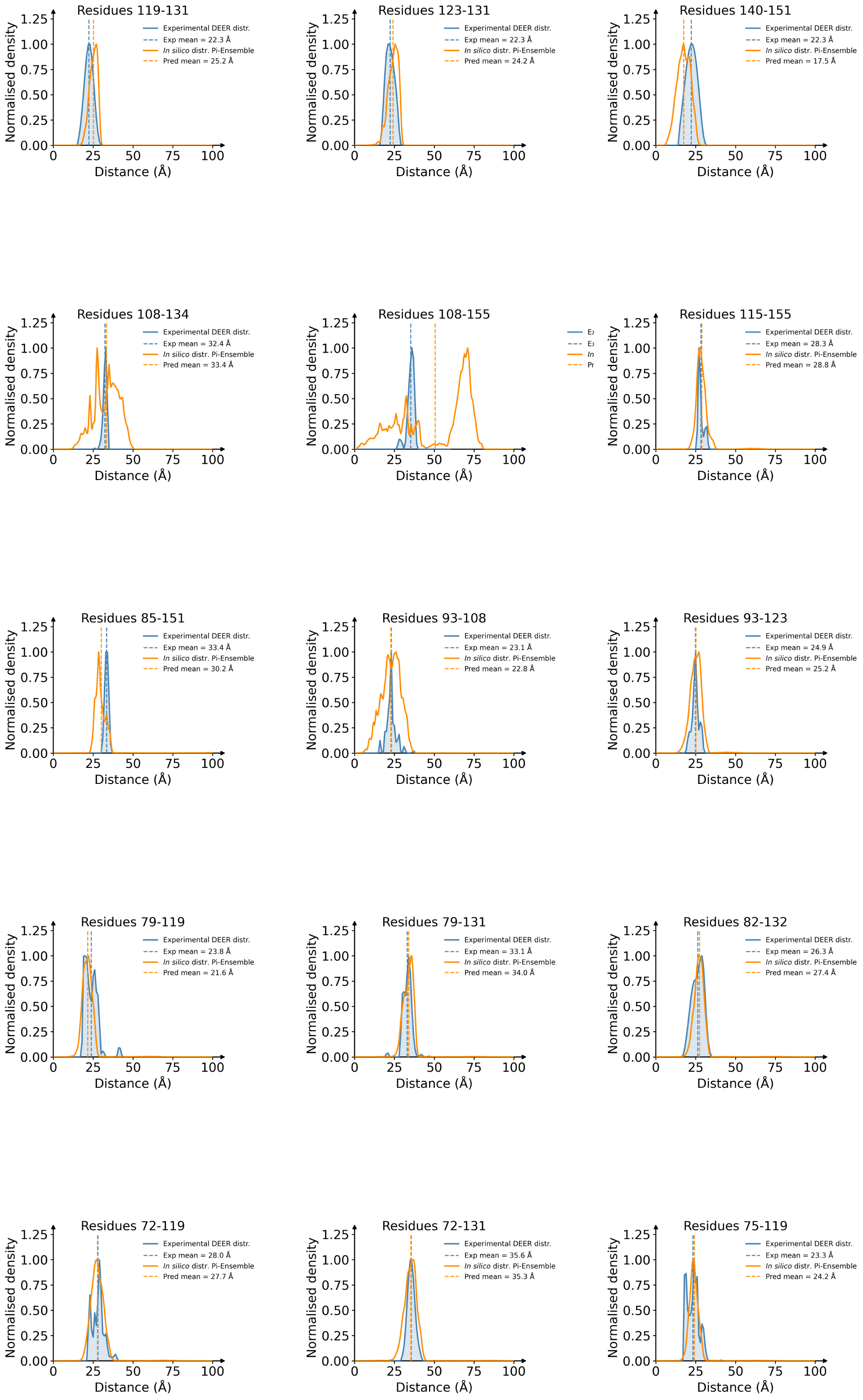


Figure S13: Overlap between experimental DEER distributions and the corresponding Pi-Ensemble predictions for T4 lysozyme residue pairs (plots 1 to 15 out of 51; continues).

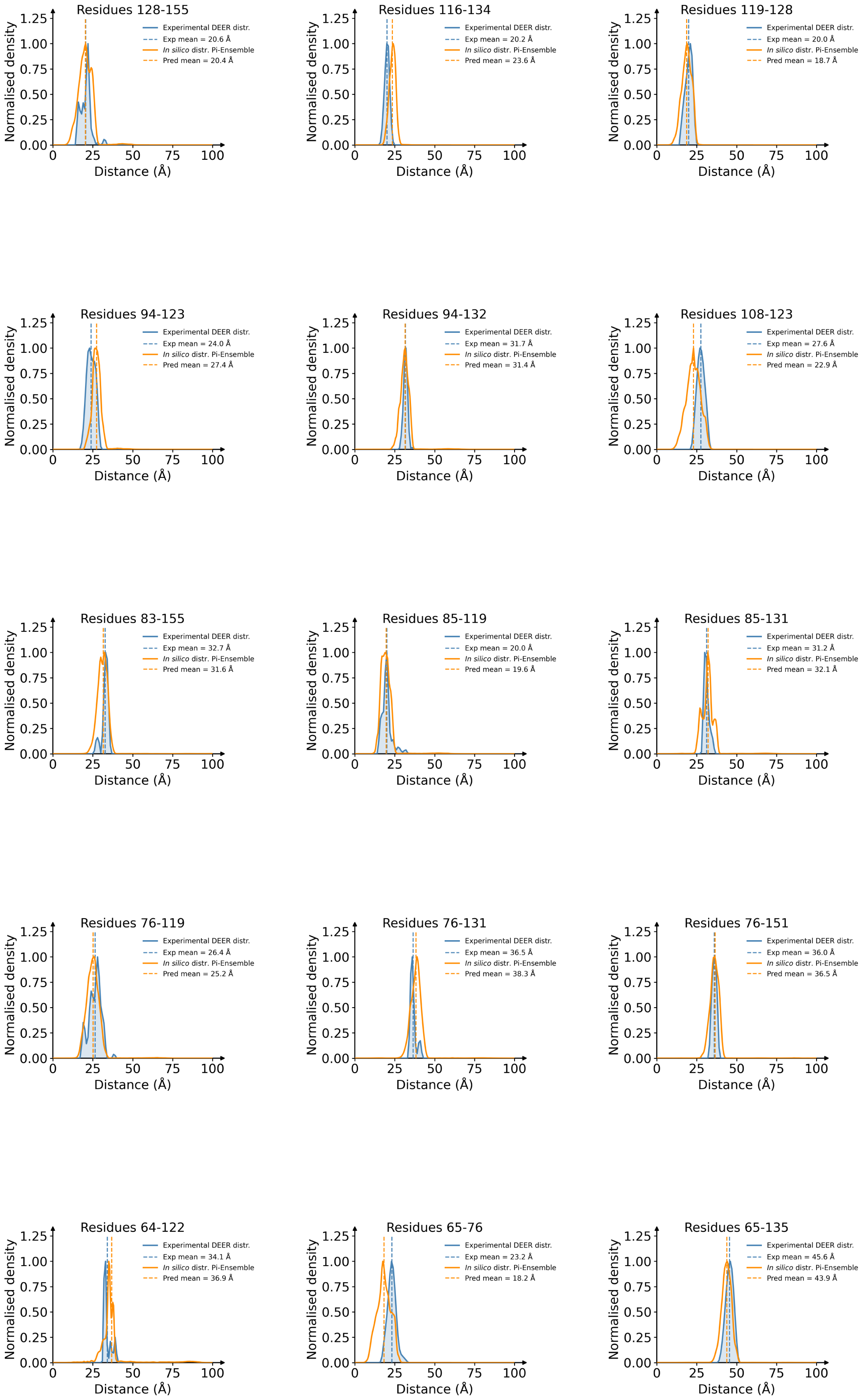


Figure S14: Overlap between experimental DEER distributions and the corresponding Pi-Ensemble predictions for T4 lysozyme residue pairs (plots 16 to 30 out of 51; continues).

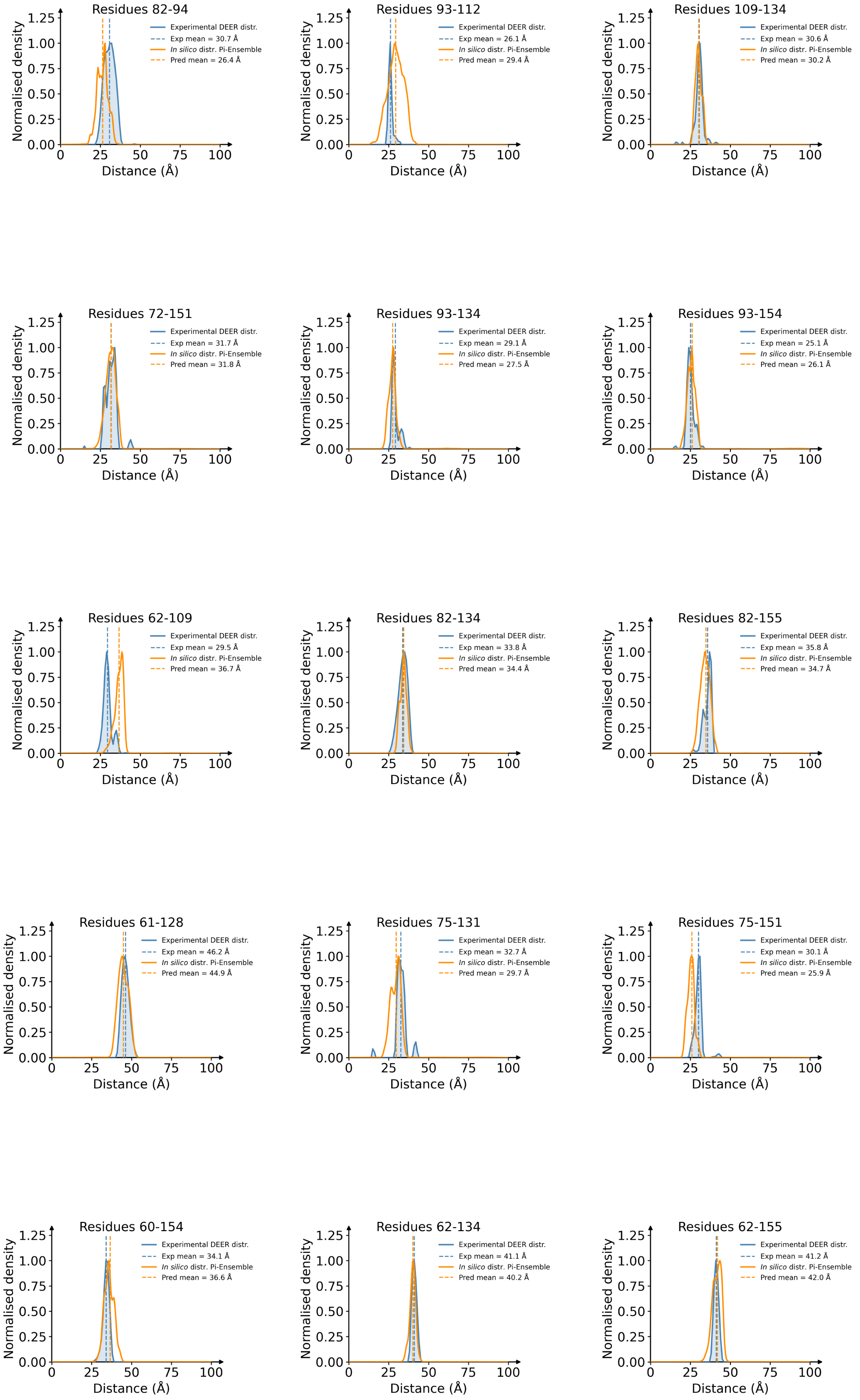


Figure S15: Overlap between experimental DEER distributions and the corresponding Pi-Ensemble predictions for T4 lysozyme residue pairs (plots 31 to 45 out of 51; continues).

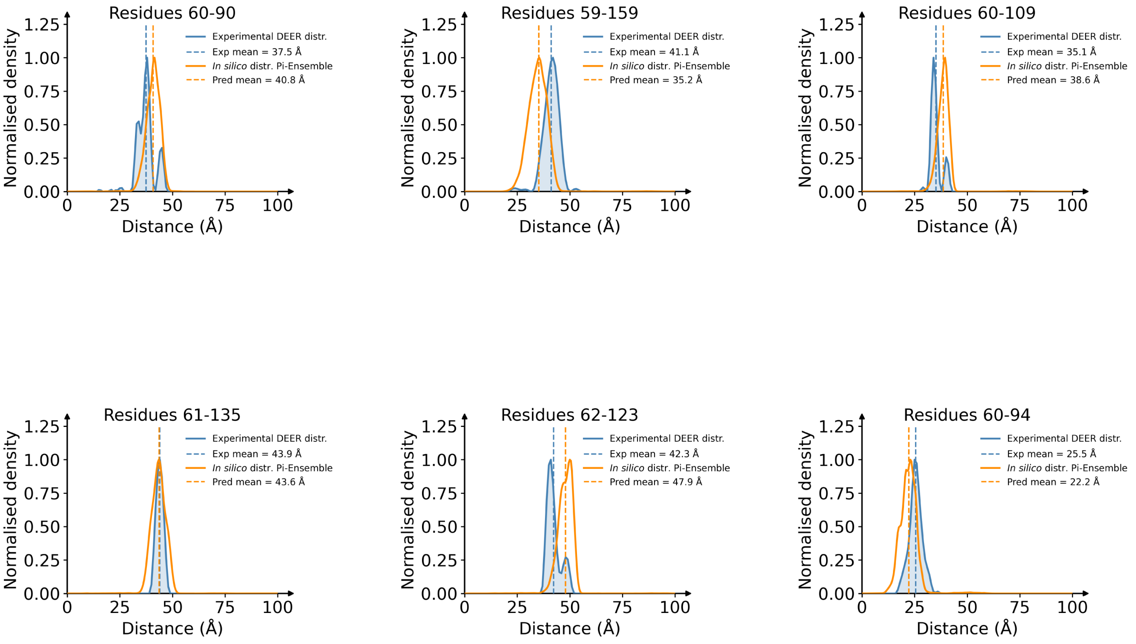


Figure S16: Overlap between experimental DEER distributions and the corresponding Pi-Ensemble predictions for T4 lysozyme residue pairs (plots 46 to 51 out of 51).
